# A Model-Driven Meta-Analysis Supports the Emerging Consensus View that Inhibitory Neurons Dominate BOLD-fMRI Responses

**DOI:** 10.1101/2024.10.15.618416

**Authors:** Nicolas Sundqvist, Henrik Podéus, Sebastian Sten, Maria Engström, Salvador Dura-Bernal, Gunnar Cedersund

## Abstract

Functional magnetic resonance imaging (fMRI) is a pivotal tool for mapping neuronal activity in the brain. Traditionally, the observed hemodynamic changes are assumed to reflect the activity of the most common neuronal type: excitatory neurons. In contrast, recent experiments, using optogenetic techniques, suggest that the fMRI-signal instead reflects the activity of inhibitory interneurons. However, these data paint a complex picture, with numerous regulatory interactions, and where the different experiments display many qualitative differences. It is therefore not trivial how to quantify the relative contributions of the different cell types and to combine all observations into a unified theory. To address this, we present a new model-driven meta-analysis, which provides a unified and quantitative explanation for all data. This model-driven analysis allows for quantification of the relative contribution of different cell types: the contribution to the BOLD-signal from the excitatory cells is <20 % and 50-80 % comes from the interneurons. Our analysis also provides a mechanistic explanation for the observed experiment-to-experiment differences, e.g. a biphasic vascular response dependent on different stimulation intensities and an emerging secondary post-stimulation peak during longer stimulations. In summary, our study provides a new, emerging consensus-view supporting the larger role of interneurons in fMRI.

## Introduction

Functional magnetic resonance imaging (fMRI) is a cornerstone of neuroimaging, as it allows researchers and clinicians to assess neuronal activation in response to stimuli (Figure 1A)(Gore, 2003; Kim and Ogawa, 2012). fMRI does not measure neuronal activity directly but is instead sensitive to changes in the ratio of oxygenated and deoxygenated blood (Ogawa *et al*., 1990; Kim and Ogawa, 2012). Thus, the neuronal activity determined from fMRI reflects temporal changes in the blood oxygenation levels i.e., the so-called Blood Oxygen Level-Dependent (BOLD) response (Ogawa *et al*., 1990; Hillman, 2014). This fMRI-BOLD response relies on a balance between the cerebral metabolic rate of oxygen (CMRO_2_) and the active regulation of the cerebral blood flow (CBF), both of which are coupled to the neuronal activity via the neurovascular coupling (NVC)(Figure 1B)(Logothetis, 2003; Hillman, 2014). The conventional interpretation of the fMRI-BOLD signal generally assumes that changes in the BOLD signal echo increases and decreases in excitatory, i.e. stimulating, neuronal activity (Logothetis *et al*., 2001; Logothetis, 2008). In other words, the general assumption is that regions with excitatory activation upregulate the CBF, which leads to an increased fMRI-BOLD signal. In the same way, the assumption has been that activation of inhibitory neurons should inhibit further activity in nearby neurons, leading to a decrease in the fMRI-BOLD signal.

**Figure 1:**
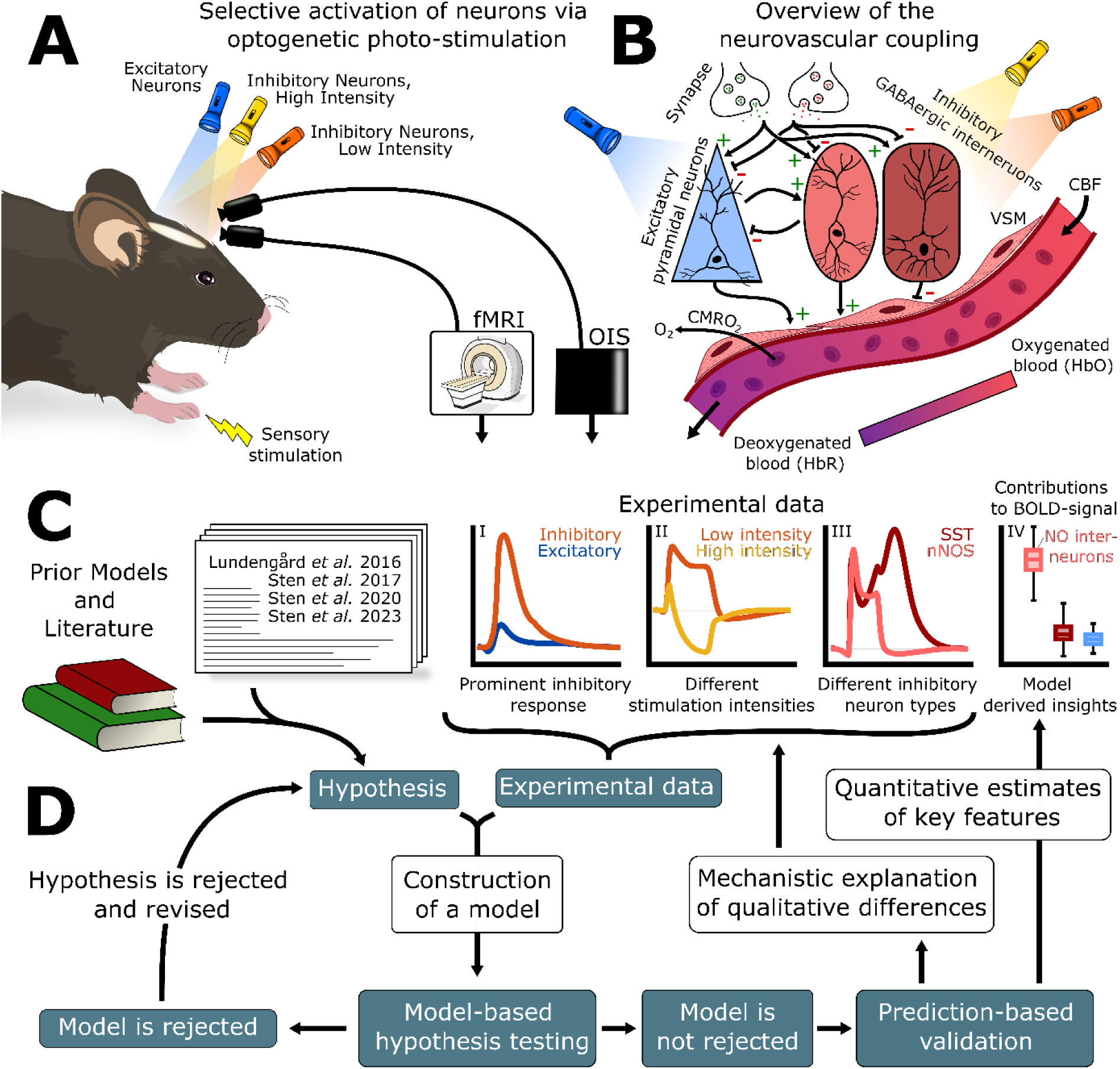
An overview of the study. **A)** Highly detailed data has been collected from mice, using optogenetic stimulation of inhibitory (orange and yellow) and excitatory (blue) neurons, as well as sensory stimulation. These data measure the changes in the blood oxygen saturation, either using fMRI-BOLD or optical intrinsic signalling (OIS) imaging. **B)** Overview of the neurovascular coupling (NVC). Neurotransmitters are released from the synaptic cleft, which either upregulate or downregulate surrounding neurons. Excitatory pyramidal neurons excite surrounding neurons and inhibitory GABAergic interneurons inhibit surrounding neuronal activity. The process of releasing neurotransmitters is energy-intensive, and the cerebral metabolic rate of oxygen (CMRO_2_) is high. To replenish metabolites, some neurons can dynamically control the dilation and constriction of cerebral blood vessels through the release of vasoactive substances that act on the vascular smooth muscles (VSM) surrounding the blood vessel. **C)** These highly detailed data show: I) differences between inhibitory and excitatory stimulations (orange vs blue), II) stimulation intensity dependencies between low-intensity (orange line) and high-intensity (yellow line), and III) different response profiles when inhibitory stimulation is applied to the different GABAergic interneurons (light red-NO and crimson-SST). **D)** These different attributes in data can be analysed using mathematical modelling. A hypothesis is formulated as a model and evaluated towards the experimental data. The hypothesis is rejected if it cannot describe the data, which leads to revision of the hypothesis. This is repeated until a hypothesis that cannot be rejected is obtained. Using this unrejected model, prediction-based model validation is obtained when the model can predict data it has not been trained to. Finally, such a validated model is used to obtain mechanistic explanations of qualitative differences (I-III) and to estimate and quantify key features (IV). Herein, this approach is used to show that NO-interneurons (IV) are the biggest contributors to the BOLD response.

Contrary to this traditional interpretation, several recent studies (Vazquez *et al*., 2014; Uhlirova *et al*., 2016; Vazquez, Fukuda and Kim, 2018; Desjardins *et al*., 2019; Echagarruga *et al*., 2020; Krawchuk *et al*., 2020; Lee *et al*., 2020; Moon *et al*., 2021) show that *γ*-aminobutyric acid (GABA)ergic inhibitory interneuron populations affect the regulation of CBF and blood oxygen saturation to a larger extent than excitatory neurons (Figure 1C, I). These studies typically use optogenetics, i.e. genetically modified mice to with a light pulse selectively activate either excitatory or inhibitory neurons (Figure 1A). Following the stimulation, hemodynamic responses, such as CBF and BOLD-signal responses, are measured using e.g. fMRI or optical imaging of the optical intrinsic signal (OIS). These measurements generally imply that the relationship between the BOLD response and neuronal activity is more complicated than conventionally assumed. This is in part due to the high complexity of the involved regulatory systems, which makes it hard to quantify the contribution of the different cell types to the NVC. Furthermore, the differences in the experimental conditions of these studies mean that their findings display numerous qualitative and quantitative differences in the observed responses (e.g. Figure 1C, I-III). In summary, as the mechanisms behind these qualitative differences are not fully understood, obtaining a unified explanation for all these responses is challenging (Figure 1C, IV).

An approach to establishing such a unified explanation is mathematical modelling. Mathematical modelling allows for integration of prior understanding with newer experimental findings and hypotheses, to gain a more holistic representation of our mechanistic understanding of the NVC (Figure 1D). There have been previous model efforts that have investigated NVC mechanisms (Griffeth and Buxton, 2011; Havlicek *et al*., 2015; Kim and Ress, 2016; Lundengård *et al*., 2016; Di Volo *et al*., 2019; Sten *et al*., 2020, 2023; Moon *et al*., 2021; Tesler, Linne and Destexhe, 2023). However, no existing model can provide a mechanistic explanation for the new optogenetics data (Vazquez, Fukuda and Kim, 2018; Lee *et al*., 2020; Moon *et al*., 2021), that indicates a larger role for interneurons.

In this work, we present a mathematical framework that: i) quantifies cell-type-specific contributions to the fMRI-BOLD signal and ii) can explain the qualitative differences in the vasoactive response across multiple experiments. By aggregating and simultaneously analysing data from multiple studies, this framework allows us to establish a model-driven meta-analysis of the mechanisms making up the NVC. This approach presents a critical step towards forming a consensus view of how inhibitory neurons contribute to the NVC, a view which contradicts the traditional interpretation of the fMRI-BOLD signal. As such, the qualitative and quantitative explanations presented in this work argue in support of a paradigm shift in how neuroimaging can, and cannot, be used to infer neuronal activity.

## Results

We here present a mechanistic, model-driven, meta-analysis that supports the emerging consensus-view that inhibitory neurons dominate the vascular regulation of the NVC. Our analysis offers: i) quantitative estimates of the cell-type-specific contributions to the neurovascular response (Figure 2) and ii) mechanistic explanations of the qualitative differences between observed responses across different studies (Figures 3-7). More specifically, the quantitative estimations of the cell-type-specific contribution to the BOLD signal are based on a mechanistic model trained on data from 11 different experiments (Figures 2A and D). These results show that the inhibitory neurons contribute >50 % of the BOLD signal in 8 of the 11 experiments and >75 % of the vascular response in all experiments (Figure 2B).

**Figure 2:**
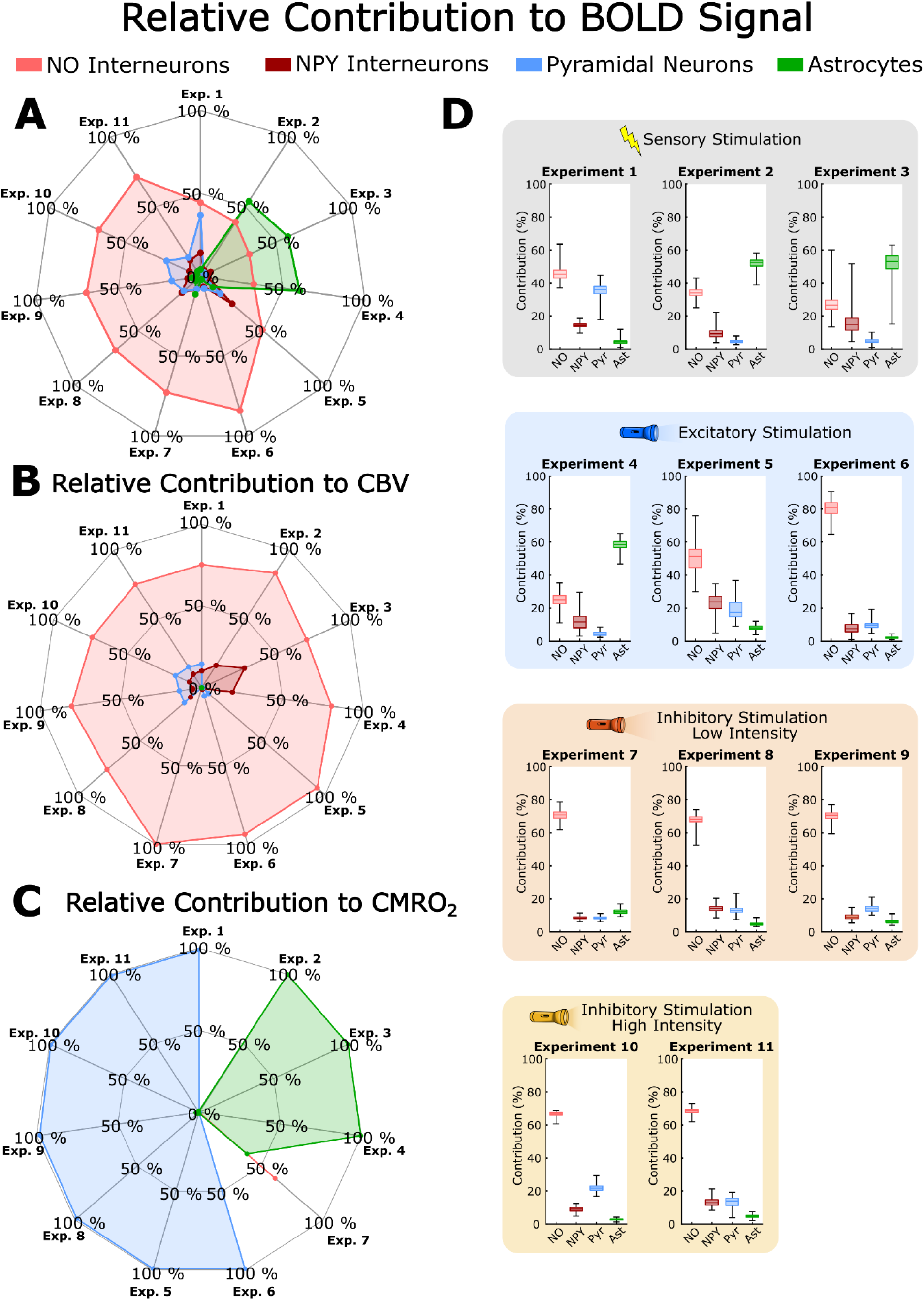
The contribution of different neuronal populations to the BOLD signal, available thanks to our model-based analysis. **A)** The contribution of four different neural populations: nitric oxide (NO) interneurons (light red), neuropeptide Y (NPY) interneurons (crimson), pyramidal neurons (Pyr, blue), and astrocytes (Ast, green), to the blood oxygen level-dependent (BOLD) signal are presented for 11 different experiments. **B)** The contributions of the four neurons (NO, NPY, Pyr, and Ast) to cerebral blood volume (CBV) response in the 11 experiments. **C)** The contributions of the four neurons (NO, NPY, Pyr, and Ast) to cerebral metabolic rate of oxygen (CMRO_2_) response in the 11 experiments. **D)** Boxplots detailing the contribution of the four neuronal populations (NO light red, NPY crimson, Pyr blue, Ast green), with a 95 % confidence interval, for the 11 experiments presented in A. The background of each experiment indicates the type of stimulation that generated these behaviours: grey is a sensory stimulation, blue is an optogenetic excitatory stimulation, orange is an optogenetic inhibitory low-intensity stimulation, and yellow is an optogenetic inhibitory high-intensity stimulation.

**Figure 3:**
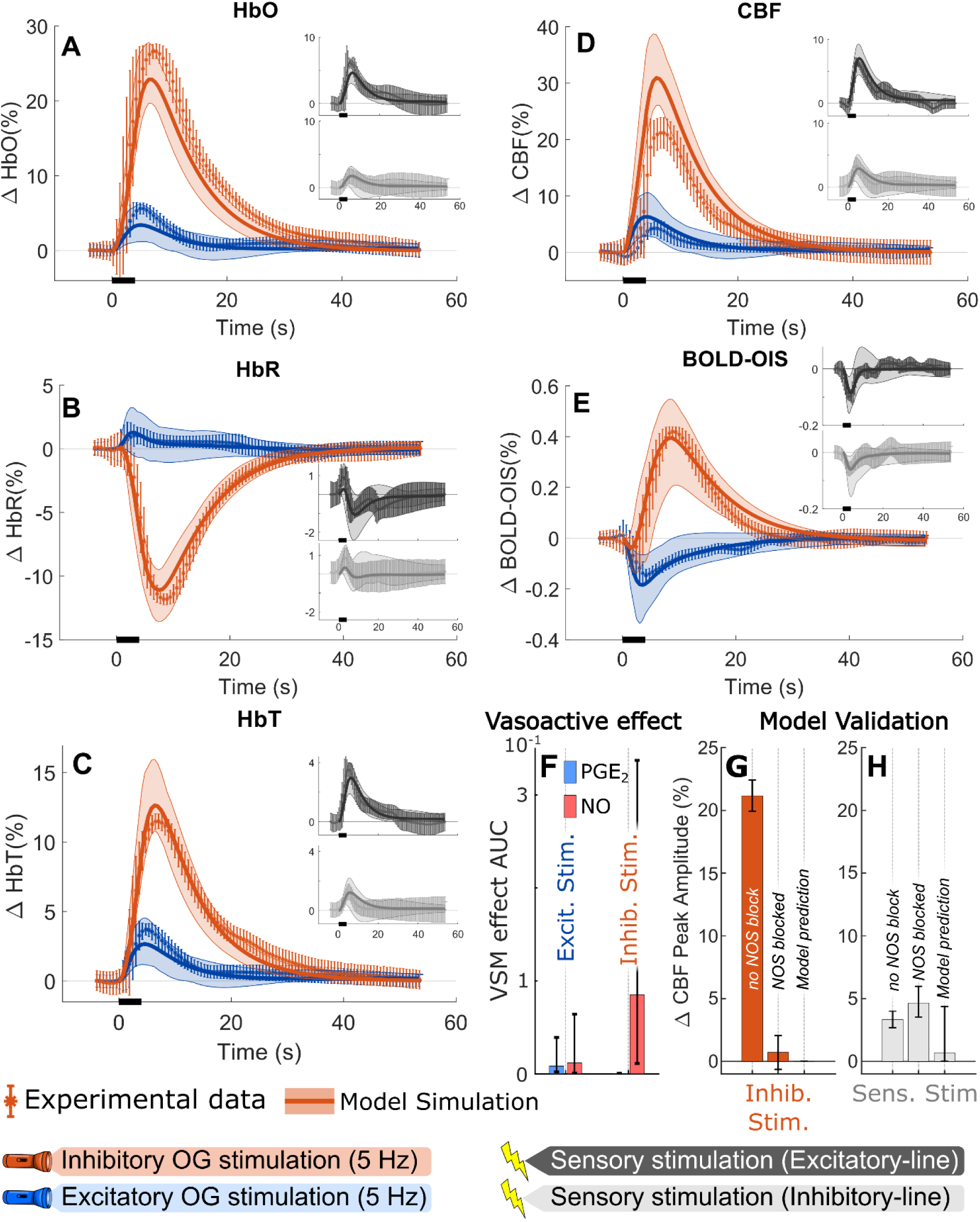
The model can describe a neurovascular response driven by the inhibitory neurons. Data and model simulations for inhibitory optogenetic 5 Hz stimulation (orange), excitatory optogenetic 5 Hz stimulation (blue), sensory electrical stimulation on the inhibitory mice-line (grey), and sensory electrical stimulation on the excitatory mice-line (dark grey). The stimulation duration is indicated by the black bar at the x-axis and is 4 s long (**A-E, G, H**). Data is originally presented by Vazquez et al. (Vazquez, Fukuda and Kim, 2018) and describe percentual concentration changes (Δ%) of **A)** oxygenated haemoglobin (HbO), **B)** deoxygenated haemoglobin (HbR), **C)** total haemoglobin (HbT), **D, G-H)** cerebral blood flow (CBF), and **E)** the OIS-BOLD signal. For each graph: the mean experimental data is indicated by an asterisk (*); the standard error of the mean is given by the error bars; the best model simulation is displayed as the solid line; the model uncertainty is given by the coloured semi-transparent overlay, corresponding to a 95 % CI. The sensory data and simulation are shown in the inserted graphs (A-E). The x-axis represents time in seconds. **F**) The model simulated area under the curve (AUC) for the overall vasoactive effect of NO, inhibitory dilatating vasoactive substance, and PGE_2_, excitatory dilating vasoactive substance, are shown for the excitatory and inhibitory simulations. Error bars indicate a 95 % CI for the AUC (F-H). A model validation of blocking the NOS signalling pathway was tested by: **G)** stimulating with a 5 s long inhibitory 5 Hz stimulation and **H)** a 5 s long sensory electrical stimulation. The bars show the peak amplitude percentage change (Δ%) in cerebral blood flow. The left bar represents the measured peak amplitude when the NOS pathway is not blocked, corresponding to the data error bars in panel D. The middle bar represents the measured peak amplitude when the NOS pathway is blocked. The right bar represents the model-predicted peak amplitude when the NOS pathway is blocked.

### Quantitative cell type contributions to the BOLD signal

Our model-based meta-analysis consists of two steps: i) Develop a model that can simultaneously fit data from all considered studies (Figures 1D, 3, 5, and 7) and correctly predict new validation data (Figures 3G-H, and 4), ii) Analyse the validated model, to draw mechanistic conclusions (Figures 3, 6, and S3). Using this model-driven approach to the data analysis, we can quantitatively estimate the contribution of different neuronal populations to the BOLD-response. Compiling these estimates across the 11 different considered experiments, it is clear that inhibitory interneurons contribute more to the BOLD-response than excitatory neurons (Figure 2A and D). Figure 2A illustrates this compilation in the form of a spider plot, where the different neurons (and astrocytes) are represented by colour and their relative contributions to the BOLD-responses for each experiment are plotted along the axes. The 11 experiments can be divided into 4 different stimulation paradigms: sensory stimulation, optogenetic stimulation of excitatory neurons, low-intensity optogenetic stimulation of inhibitory neurons, and high-intensity optogenetic stimulation of inhibitory neurons. As can be seen, the combined contributions of the interneurons to the BOLD-signal are around 50-80 %, while the pyramidal neurons contribute <20 %. A more detailed breakdown of the contributions reveals that the combined contributions of NO and neuropeptide Y (NPY) interneurons (Figure 2D, light red and crimson bars) are larger than the contributions of pyramidal neurons (Figure 2D, blue bars) in all 11 experiments. Note that these results cannot be obtained by a mere inspection of the original data and that it stands in contrast to the conventional interpretation of BOLD-fMRI, which assumes that the signal comes predominantly from excitatory pyramidal cells.

**Figure 4:**
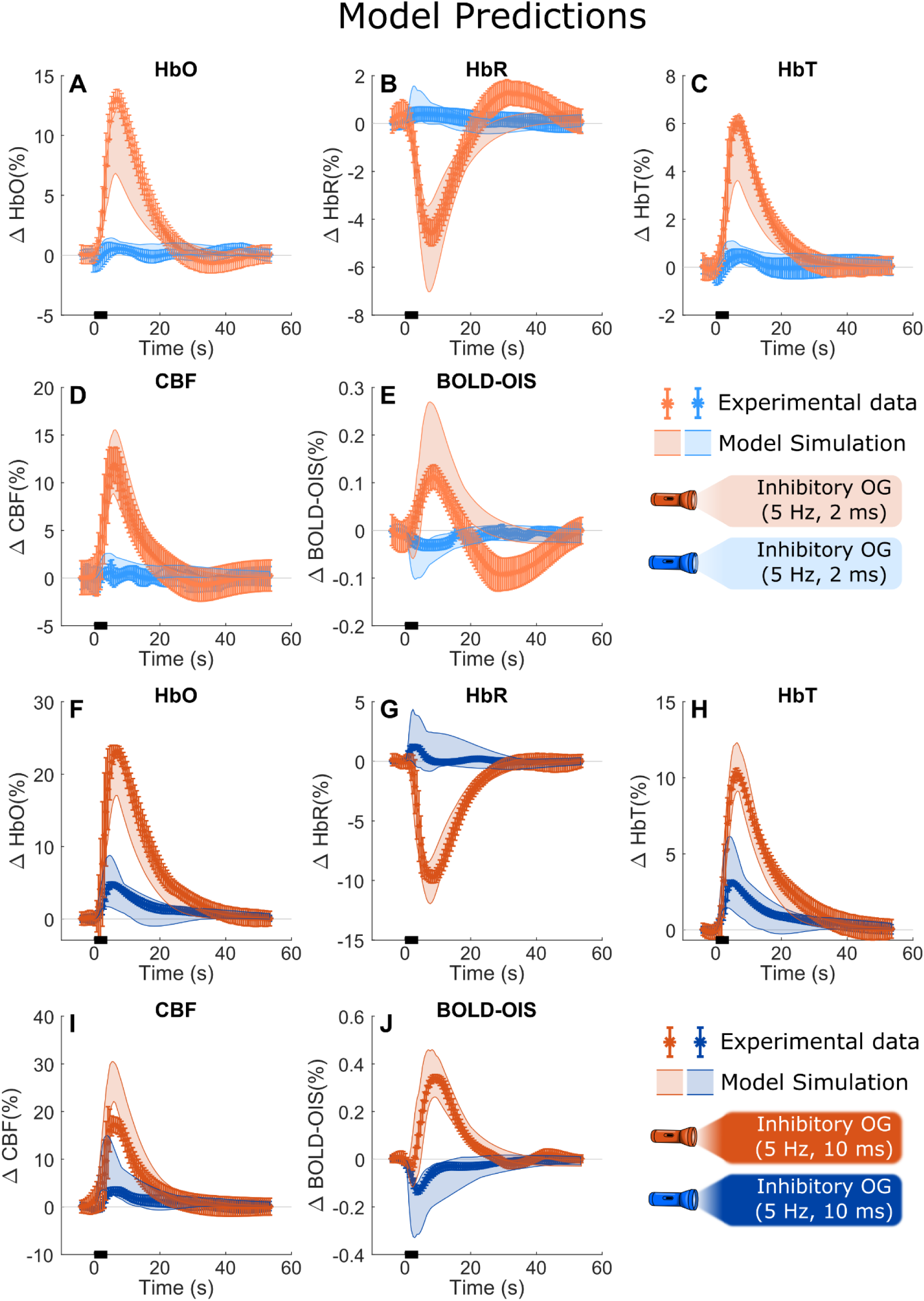
The model can predict OG data which it has not been trained on. Data and model simulations for optogenetic stimulation with a frequency of 5 HZ and the type: i) inhibitory stimulation with 2 ms pulse width (light orange), ii) excitatory stimulation with 2 ms pulse width (blue), iii) inhibitory stimulation with 10 ms pulse width (orange), and iv) excitatory stimulation with 10 ms pulse width (dark blue). The stimulation duration is indicated by the black bar at the x-axis and is 4 s long (**A-J**). Data is originally presented by Vazquez et al. (Vazquez, Fukuda and Kim, 2018)and describe percentual concentration changes (Δ%) of oxygenated haemoglobin (HbO)(**A, F**), deoxygenated haemoglobin (HbR)(**B, G**), total haemoglobin (HbT)(**C, H**), cerebral blood flow (CBF)(**D, I**), and the OIS-BOLD signal (**E, J**). For each graph: the mean experimental data is indicated by asterisk (*); the standard error of the mean is given by the error bars; the model uncertainty is given by the coloured semi-transparent overlay, corresponding to a 95 % CI.

**Figure 5:**
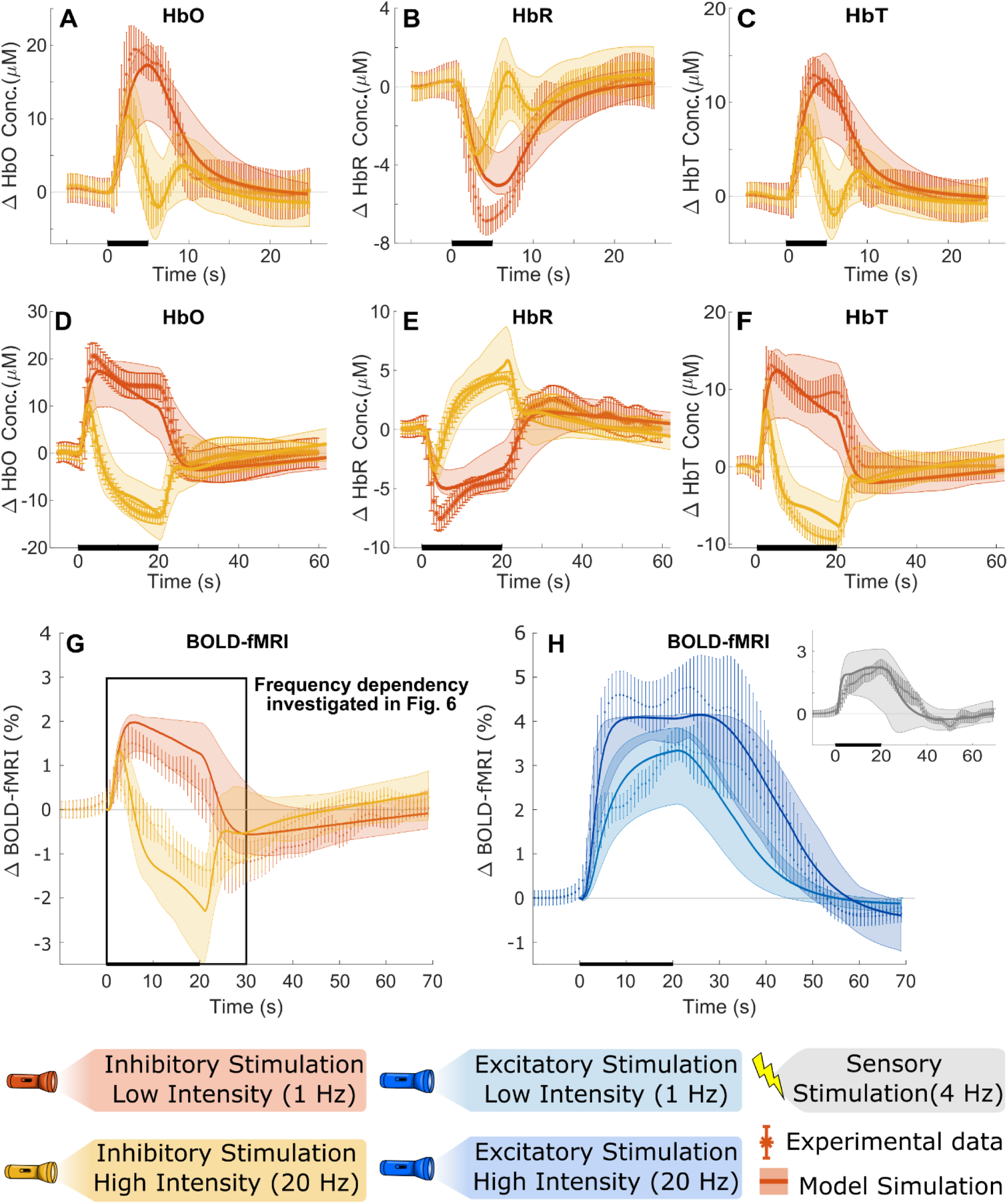
The model can simultaneously describe the qualitatively different responses to the different stimulation intensities. Data and model simulations for inhibitory optogenetic 20 Hz stimulation (yellow) and inhibitory optogenetic 1 Hz stimulation (orange). The stimulation duration is denoted by the black bar at the x-axis, 5 s long (**A-C**) and 20 s long (**D-F**). Data was originally presented by Moon et al. (Moon et al., 2021) and describe the relative concentration changes (ΔµM) of oxygenated haemoglobin (HbO)(**A, D**), deoxygenated haemoglobin (HbR)(**B, E**), and total haemoglobin (HbT)(**C, F**). Data for BOLD-fMRI for these two inhibitory stimulations is presented in **G)**. Optogenetic excitatory stimulations were also performed with the same frequencies, blue 1 Hz and dark blue 20 Hz, and the BOLD-fMRI response is shown in **H)**. The insert shows the response for a 4 Hz sensory stimulation, grey. For each graph: the mean value of the experimental data is indicated by an asterisk (*); the standard error of the mean is given by the error bars; the best model simulation is seen as the solid line; the model uncertainty, corresponding to a 95 % CI, is given by the coloured semi-transparent overlay. The x-axis represents time in seconds.

**Figure 6:**
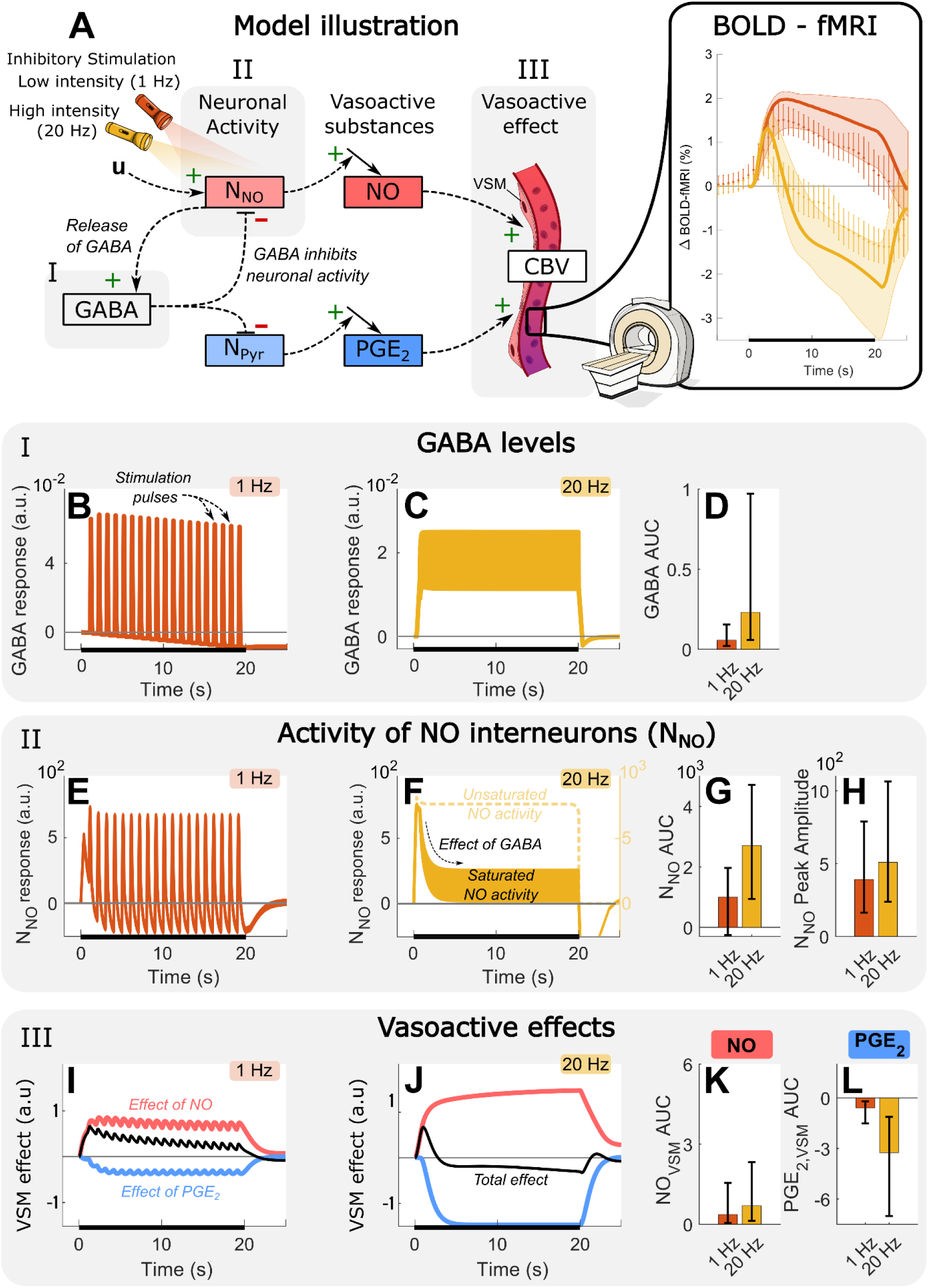
The neurovascular dependency on the stimulation intensity is explained by GABA dynamics. **A)** A simplified model illustration, showing that the stimulation (u) upregulates neuronal activity of NO interneurons (N_NO_) and pyramidal neurons (N_Pyr_). The neuronal activity then leads to production of vasoactive substances, NO and PGE_2_. Additionally, N_NO_ initiates the release of GABA, which downregulates all neuronal activity. Thus, the vasoactive response is a result of the balance between the effect of the stimulation and the inhibitory effect of GABA. The biphasic behaviour seen in the Moon et al. (Moon et al., 2021) data is showcased in the box to the right, with the 1 Hz stimulation being orange and 20 Hz stimulation being yellow. The simulated model behaviour of B) GABA for the 1 Hz and C) 20 Hz stimulation, show that D) the AUC of GABA is larger for the 20 Hz stimulations. E) the simulated model behaviour of N_NO_ for the 1Hz stimulation, and F) the 20 Hz stimulation, similarly show that the G) area under the curve (AUC) and H) peak value is relatively similar. This would not be the case if the stimulation effect was not subject to saturation, F right y-axis. The 1 Hz stimulation produces the vasoactive effect in I) where the total effect (black) reflects the dilating effect of NO (red), as the constricting effect from a lack of PGE_2_ is weaker. J) The 20 Hz stimulation results in a net vasoconstricting effect (black curve below baseline) due to a more substantial lack of PGE_2_ which cancels out the dilating effect of NO. Notably, the increase K) of NO levels is not as big compared to the loss L) of PGE_2_ comparing the high- and low-intensity stimulations. Error bars indicate a 95 % CI.

These results show that in 8 out of the 11 cases, the NO-interneurons are the largest contributors to the BOLD-signal, even in the cases of optogenetic excitatory stimulation (Figure 2D, blue box). For the three remaining cases, NO-interneurons still have a large contribution, but the model also attributes a major portion of the BOLD-signal to the astrocytes (Figure 2D, green bars). Consulting the model, we can further investigate this behaviour, by subdividing the quantitative contributions to the BOLD-response into a vascular contribution (changes in CBV, Figure 2B) and a metabolic contribution (changes in CMRO_2_, Figure 2C). Studying these sub-contributions, two results stand out: i) NO-interneurons are in all 11 cases the largest contributor to changes in CBV (Figures 2B and S1), and ii) pyramidal neurons are in 7 of 11 cases the largest contributor to changes in CMRO_2_ (Figures 2C and S1). Lastly, for the three cases where the astrocytes contributed the most to the BOLD-response (Figure 2A, green), the model attributes this behaviour to the astrocytes having a large effect on the changes in CMRO_2_ (Figures 2C and S1, green) combined with relatively small changes in the total CBV response. A detailed breakdown of how these quantitative estimates were acquired can be found in Supplementary Material (Supplementary Material section 4). Let us now turn to the second usage of the validated model: providing a mechanistic underpinning for observations, such as the high role of inhibitory neurons.

### Inhibitory neurons as a driving force for vascular regulation

The results presented above (Figure 2), are contingent on the model being an accurate explanation of the experimental data. The agreement between model and data are shown in Figures 3 and 5. Let us first consider the results in Figure 3, which focuses on the differences between inhibitory and excitatory stimulations (Vazquez, Fukuda and Kim, 2018). This data shows: the measured changes in haemoglobin (oxygenated haemoglobin, HbO, Figure 3A; reduced haemoglobin, HbR, Figure 3B; total haemoglobin, HbT, Figure 3C), the laser Doppler flowmetry (LDF) measurements of CBF (Figure 3D), and the relative percentage change in OIS measurements for BOLD and Hb (Figure 3E). All these data show the result of both an optogenetic excitatory stimulation (blue), and an optogenetic inhibitory stimulation (orange). Note that the data show a larger response in Hb levels, CBF, and OIS-BOLD for stimulation of the inhibitory neurons (Figure 3 orange). Note also that the model simulations (continuous lines and shaded areas) agree with all the main features seen in the data (error bars). For example, changes in HbT following an inhibitory stimulation (Figure 3C, orange) show a prominent peak culminating at around 12 %, at ca 10 sec, for both model simulation and data. Following this peak, the response then subsides and returns to baseline. This qualitative assessment was also confirmed using a *χ*^2^-test (Equation 11; with a test statistic *f*(*θ*_*H*1_) = 2033.43 that is lower than the cut-off threshold *χ*^2^(*α* = 0.05, *Dof* = 3347) = 3482.70).

A more in-depth analysis of the model shows that NO released from NO-interneurons is the dominant vasoactive agent. Model simulations of the two main vasodilating substances, NO and Prostaglandin E_2_ (PGE_2_), can be seen in Figure 3F, for both excitatory and inhibitory stimulation. As expected, for the inhibitory stimulation, NO is the dominant vasoactive agent (Figure 3F, right). Interestingly, also for excitatory stimulation, NO has a contribution equally big as the PGE_2_ contribution (Figure 3F, left).

The idea that NO is the primary vasoactive substance is explored by a model validation test, simulating the effects of blocking the nitric oxide synthase (NOS) pathway. The model predicts that the neurovascular response is eliminated (Figure 3G, right) when NOS is blocked, and the cells are exposed to an inhibitory optogenetic stimulation. This model prediction was then compared to data from Vazquez *et al*. where this experiment was done (Figure 3G, middle). As can be seen, the experimental data confirms the model prediction that the NOS-blocked response is almost non-existing (<3 %, Figure 3G, middle). A similar validation was done also for NOS being blocked during a sensory stimulation paradigm. Here, the model predicts that the response is slightly higher (0-5 %, Figure 3H, right) which is consistent with the experimental validation experiment (4-6 %, Figure 3H, middle). This low but non-zero response comes from the fact that the sensory stimulation primarily activates the excitatory pyramidal cells, due to the NOS-inhibition. Note that the model has not been trained to any of these NOS-inhibition experiments, thus the model predictions agreeing with these experiments serve as a validation of the model.

### The model is validated by its ability to predict new data not used for model training

The model was further validated by predicting additional data, presented by Vazquez *et al*. (Vazquez, Fukuda and Kim, 2018), which were not part of the model training and therefore unknown to the model. These model predictions, of optogenetic inhibitory and excitatory stimulation with pulse widths of 2 ms and 10 ms, are in good agreement with data and capture the main features. For instance, the amplitude of the inhibitory HbO response is accurately lowered from 23 % (Figure 4F) to 15 % (Figure 4A) for the inhibitory stimulation (orange), comparing pulse widths of 10 ms and 2 ms respectively.

### The model explains how different stimulation frequencies lead to qualitatively different vascular responses

The training and validation data used for the model-driven meta-analysis also include data showcasing a frequency-dependent behaviour in response to different optogenetic stimulations. The data, presented by Moon *et al*. (Moon *et al*., 2021), are depicted together with our model simulations in Figure 5. This data shows: changes in HbO (Figures 5A and D), HbR (Figures 5B and E), and HbT (Figures 5C and F) for a 5 s stimulation (Figures 5A-C) and a 20 s stimulation (Figures 5D-F), including both a low-intensity (1 Hz) inhibitory optogenetic stimulation (orange) and a high-intensity (20 Hz) inhibitory optogenetic stimulation (yellow). Further, the BOLD-fMRI is shown for both inhibitory stimulations (Figure 5G) and for comparison, the BOLD-fMRI response to a: low-intensity (1 Hz) excitatory optogenetic stimulation, high-intensity (20 Hz) excitatory optogenetic stimulation, and a 4 Hz sensory stimulation is shown in Figure 5H. The model simulations (continuous lines and shaded areas) agree well with all the main features seen in the data (error bars). This qualitative assessment was confirmed using a the same *χ*^2^-test as previously (*f*(*θ*_*H*1_) = 2033.43, that passed the cut-off threshold, *χ*^2^(*α* = 0.05, *Dof* = 3347) = 3482.70).

One of the interesting features seen in both model and data is a biphasic vascular response, i.e. an initial increasing above the baseline, followed by a fall below the baseline (Figure 5G and 6A, yellow). This behaviour is only seen for the high-intensity stimulation, and not seen in the low-intensity stimulation. This difference is not intuitive, as one would generally associate a more intensive stimulation with a stronger and more positive vascular response, compared to the low-intensity response. It is therefore interesting to use the model to understand how this difference can be mechanistically understood.

The model explains the biphasic response observed for the high-intensity stimulation as a balance between a saturated inhibitory neuronal activity, and elevated levels of GABA. The optogenetic stimulation (*u*) excites the NO-interneurons and GABA is released, which in turn inhibits all neuronal activity (Figure 6A, N_NO_ and N_Pyr_). For the high-intensity stimulation, there is not enough time between stimulation pulses for the GABA levels to return to baseline (Figure 6C), resulting in overall elevated levels of GABA (Figure 6D), compared to the low-intensity stimulation (Figure 6B). Simultaneously, the activity of the NO-interneurons (N_NO_) is subjected to a saturation effect (Equation 7) which, together with the elevated GABA levels, results in the overall neuronal activity being very similar between the high- and low-intensity stimulations (Figure 6G, H). Although the bursting of neuronal activity settles at a lower amplitude for the high-intensity stimulation (due to the elevated GABA levels), the resting activity between spikes does not fall below the baseline (Figure 6F). Consequently, the stimulation intensity has no discernible effect on the peak amplitude of N_NO_ (Figure 6H) and only a slight effect on the overall levels of neuronal activity (Figure 6G). Subsequently, the vasodilating effect of the NO-release remain similar between the stimulations. This can be seen both in the amplitude of NO’s vasoactive effect (Figure 6I-J, red) and the overall AUC of NO’s vasoactive effect (Figure 6K). As for the pyramidal neurons, their activity (N_Pyr_) is inhibited by GABA (Figure 6A), causing a decreased release of vasodilating PGE_2_ (Figures 6I-J, blue). Due to the elevated levels of GABA during the high-intensity stimulation, the vasoactive effect of PGE_2_ is reduced substantially more (Figure 6J, blue), compared to the low-intensity stimulation (Figure 6I, blue). Since both NO and PGE_2_ have vasodilating properties, the comparatively greater reduction of PGE_2_ (Figure 6L), compared to the modest increase of NO (Figure 6K), causes a net contraction of the blood vessels after an initial dilation (Figure 6J, black). This dynamic results in the biphasic response seen for the high-intensity stimulation (Figure 6A, yellow). The biphasic response is not present during the low-intensity stimulation due to the increase of NO offsetting the reduced levels of PGE_2_, resulting in a net dilation of the blood vessels (Figure 6A, orange).

It should be noted that the saturation of neuronal activity seems to be a crucial mechanism, without which the model is not able to describe the biphasic response accurately. In the model, this saturation is implemented by limiting the degree to which the optogenetic stimulation can stimulate the model state that represents the neuronal activity. This means that during high-intensity stimulation this model state will reach a stable plateau and additional stimulation will not increase the neuronal activity further (Figure 6F, see Methods for details). Without this saturation effect, the peak amplitude of N_NO_ would increase by an order of magnitude during the high-intensity stimulation (Figure 6F, right y-axis), resulting in much more NO being released and an increased vasodilating effect. This increased vasodilation would counteract the decreased levels of PGE_2_, resulting in net vasodilation, and the biphasic response would not be observed.

The biphasic behaviour is not present when observing a similar setup using excitatory optogenetic stimulation (Figure 5H). In this case, we see the high-intensity (dark blue) stimulation produces a faster and stronger response compared to the low-intensity (blue) stimulation. While this scenario seems similar to the inhibitory stimulation at first glance, the underlying interactions differ. Activating the pyramidal neurons release glutamate which subsequently activate the interneurons and triggers the release of GABA. Crucially, this means that a balance between excitatory and inhibitory neuronal activity is achieved for both the high- and low-intensity excitatory stimulations. This alters the tug-of-war between neuronal activation and inhibition that we saw for the inhibitory stimulation, and the vascular response instead intuitively scales in proportion to the stimulation intensity. Note that the simulation of neuronal activity is still subject to the saturation effect discussed above.

### The role of Somatostatin releasing neurons in the NVC

Finally, we investigate the role of somatostatin (SOM)-releasing interneurons in the neurovascular response, using an expanded model structure which also included SOM-interneurons. This analysis was based on data presented by Lee *et. al.* (Lee *et al*., 2020) and includes: the relative changes in HbO (Figures 7A and D), HbR (Figures 7B and E), and HbT concentrations (Figures 7C and F) in response to 20 Hz optogenetic stimulations of SOM releasing interneurons (Figure 7, crimson), of NO-interneurons (Figure 7, light red), as well as sensory stimulation (Figure 7, grey). Two stimulation lengths were evaluated, 16 sec (Figures 7A-C) and 2 sec (Figures 7 D-F). The model simulations (lines) agree with all the main features seen in the data (error bars). The qualitative assessment was again confirmed using a *χ*^2^-test (Equation 11; *f*(*θ*_*H*1_) = 1013.46 which is lower than the cut-off threshold *χ*^2^(*α* = 0.05, *Dof* = 1548) = 1640.65.

**Figure 7:**
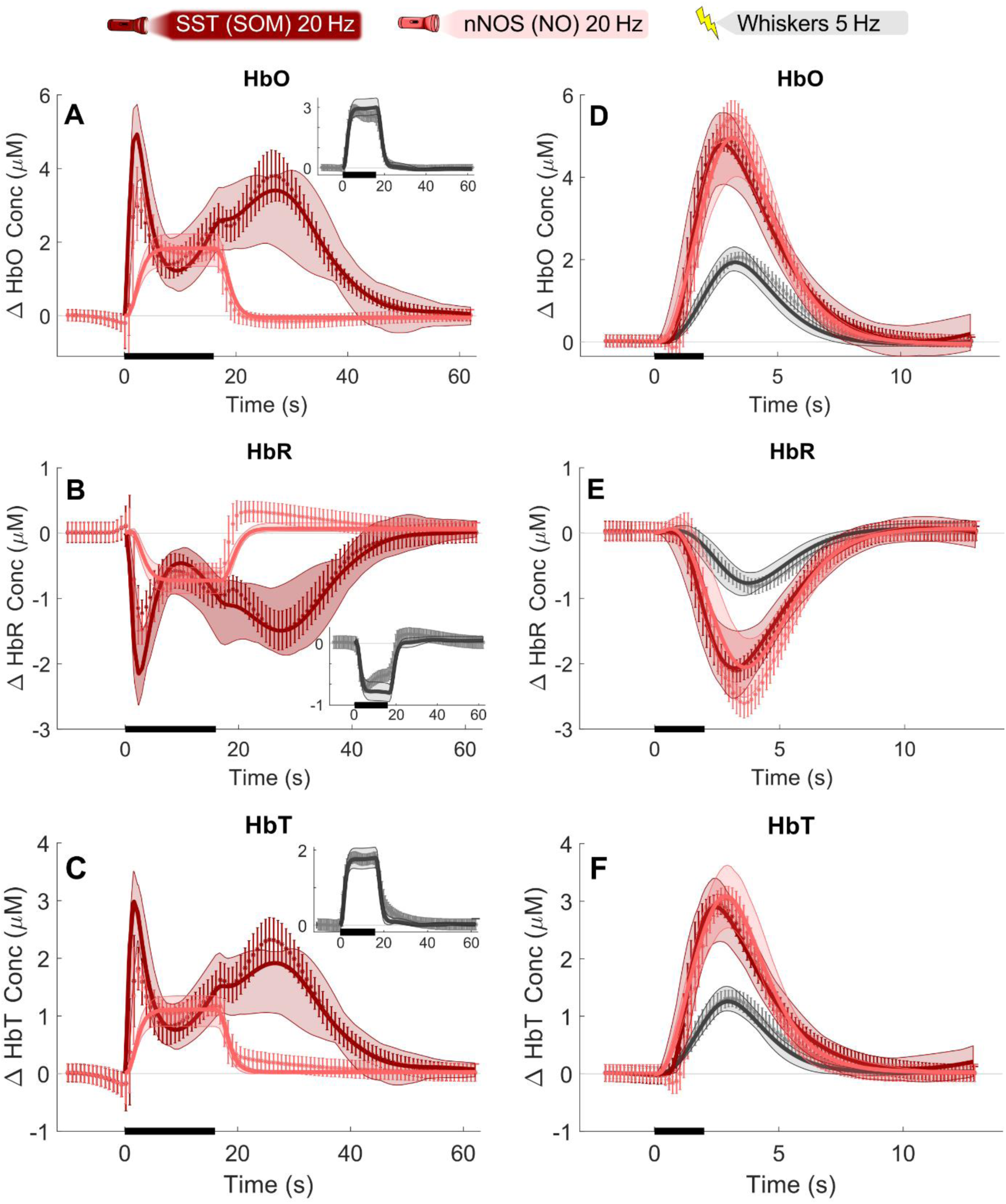
The model can describe a delayed dilation using somatostatin-releasing interneurons. Data and model simulations for inhibitory nNOS 20 Hz stimulation (light red), inhibitory SST 20 Hz stimulation (crimson), and sensory whiskers 5 Hz stimulation (grey). The stimulation duration is denoted by the black bar at the x-axis and is 16 s long (**A-C**) and 2 s long (**D-F**). Data was originally presented by Lee et al. (Lee et al., 2020) and describe the relative concentration changes (ΔµM) of oxygenated haemoglobin (HbO) (**A, D**), deoxygenated haemoglobin (HbR) (**B, E**), and total haemoglobin (HbT) (**C, F**). For each graph: the mean value of the experimental data is indicated by an asterisk (*); the standard error of the mean is given by the error bars; the best model simulation is seen as the solid line; the model uncertainty corresponding to a 95 % CI is given by the coloured semi-transparent overlay. The sensory data and simulation are shown in the inserted graphs for the long stimulation (A-C). The x-axis represents time in seconds.

The data show that selectively stimulating SOM-interneurons gives rise to a double-peak response in the Hb measurements (Figures 7A-C, crimson), which is not present for the NO-stimulation (light red), nor for the sensory stimulation (Figures 7A–C, inserts). The model can accurately describe the observed double peak dynamic in response to stimulating SOM neurons, given that a corresponding signalling pathway is introduced to the model structure (Figure 8). We found that the mechanism of this double-peak dynamic has two main components: a) the initial peak is generated by the direct vasoactive influence of the SOM-releasing neurons; b) the second peak is generated by a delayed recovery from the inhibition of activity in the other neuron populations (NO, NPY, and Pyramidal neurons). A detailed analysis of this double peak response can be found in the Supplementary Materials (Supplementary Materials section 2).

**Figure 8:**
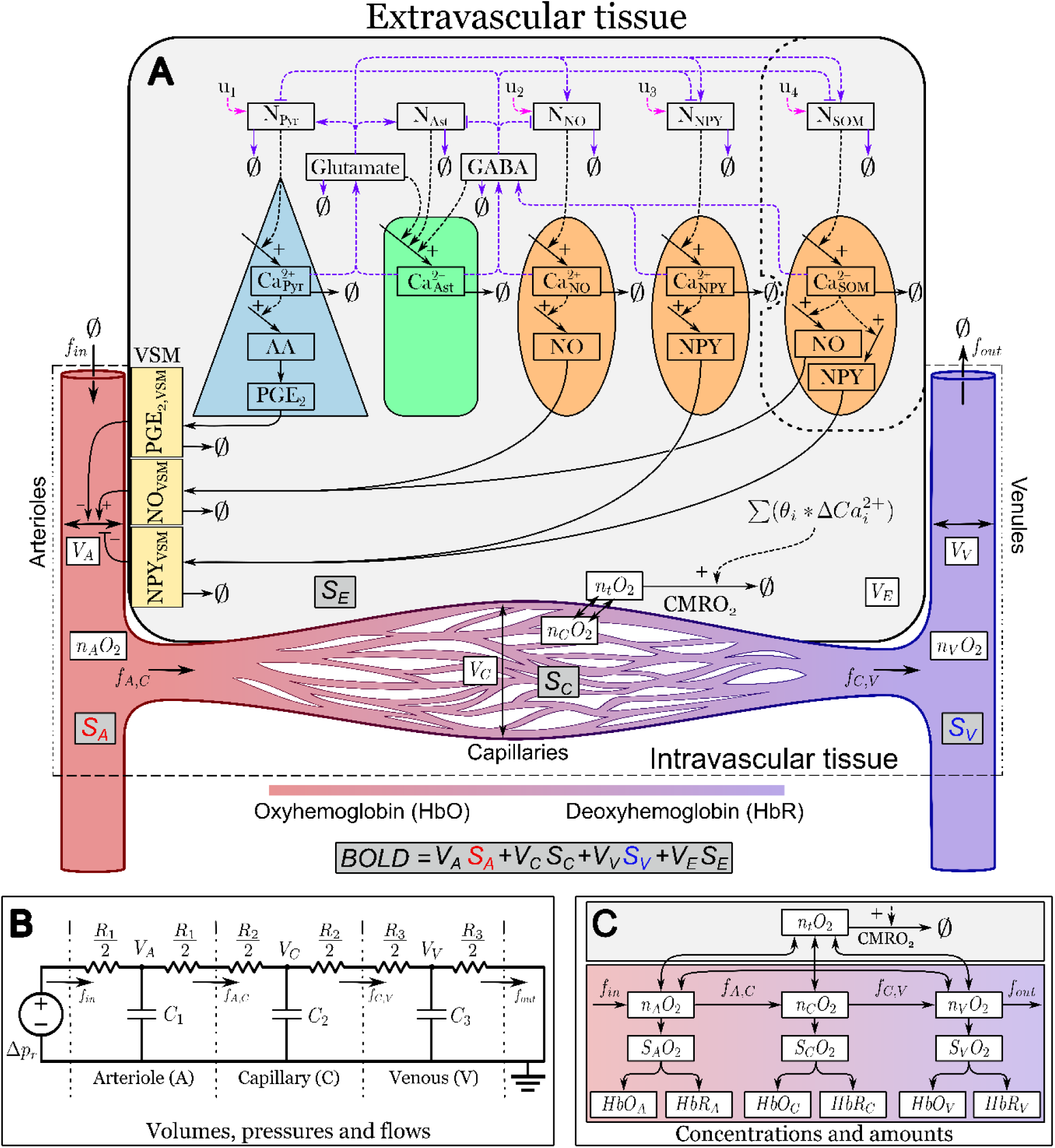
An interaction graph of the model structure. **A)** the model represents different types of neurons and glia cells, excitatory pyramidal neurons (blue), activity-regulating astrocytes (green), inhibitory GABAergic nitric oxide (NO), neuropeptide-Y (NPY), and somatostatin (SOM) releasing neurons (orange). These neurons, represented by the states *N*_*Pyr*_, *N*_*NO*_, *N*_*NPY*_, and *N*_*SOM*_, can be excited with an external stimulation, *u*_*i*={1,2,3,4}_. The excitation is unique to every unique stimulation configuration, indicated by the pink-coloured arrows. When excited the pyramidal neurons increase the level of intracellular calcium ions 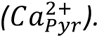 The calcium level controls the release of neurotransmitters and the activity of intracellular synthesis of vasoactive substances. The neurotransmitters released are excitatory glutamate, by pyramidal neurons, and inhibitory GABA, by GABAergic interneurons. These neurotransmitters act on the other neurons and are regulated by the astrocytes, which can both synthesize glutamate and GABA dependent on their level of intracellular calcium ions 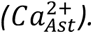 The purple-coloured arrows indicate that these signalling pathways are allowed to vary due to the anaesthesia conditions differing between different studies. The intracellular calcium is controlled by the neuronal activity of the neurons. The intracellular pathways of the excitatory and inhibitory neurons end with the release of vasoactive substances that influence the vascular smooth muscles (*VSM*) surrounding the arterioles. Pyramidal neurons release the vasodilating substance Prostaglandin E2 (PGE_2_), NO neurons release the vasodilating substance NO, NPY neurons release the vasoconstricting substance NPY, and SOM neurons release both the vasodilating substance NO and vasoconstricting substance NPY. The dilation and constriction of the arterioles control the rate of blood entering the arterioles. A generalized expression of removal or breakdown is denoted with ∅. Further, the vascular compartment of the model describes the volume (*V*_*i*={*A*,*C*,*V*,*E*}_), blood flow (*f*_*i*={*in*,*AC*,*CV*,*out*}_), oxygen content (*n*_*i*_*O*_2_, *i* = {*a*, *c*, *v*, *t*}), and cerebral metabolic rate of oxygen (CMRO2) of the arterioles (A), capillaries (C), venules (V), tissue (t), and extravascular volume (E). The compartmental volume and oxygen saturation are used to calculate the functional magnetic resonance imaging blood oxygen-level dependent (fMRI-BOLD) signal. The total fMRI-BOLD signal is the sum of the signal from each compartment (*S*_*i*={*A*,*C*,*V*,*E*}_) **B)** The three-compartment vascular model, consisting of arterioles (A), capillaries (C), and venules (V), describes the blood volumes (*V*_*i*_), pressures, and flows (*f*_*i*_) corresponding to an analogue electrical circuit. The drop in blood pressure over a compartment can be represented by a voltage drop, the blood flow is represented by an electric current, and the blood volume can be seen as an electric charge stored in capacitors. The vessel compliance (*C*_*i*={1,2,3}_) is represented by the capacitance of the capacitors, and the vessel resistance (*R*_*i*={1,2,3}_) is analogous to the electric resistance. The blood pressure difference (*ΔP*_*r*_) maintained by the intracranial pressure corresponds to the electromotive force created by a voltage source. **C)** The oxygen transport model depicts the amount of oxygen (*n*_*i*_*O*_2_,), oxygen saturation (*S*_*i*_*O*_2_, *i* = {*A*, *C*, *V*}), oxygenated haemoglobin (*HbO*_*i*={*A*,*C*,*V*}_) and deoxygenated haemoglobin (*HbR*_*i*={*A*,*C*,*V*}_), for each respective compartment. The rate of oxygen metabolism, which occurs in the extravascular tissue, is regulated by the activity of intracellular Ca^2+^ transportation in the different neurons (pyramidal, astrocyte, NO-, NPY-, SOM-interneurons). Oxygen diffusion from the vessel compartments to the extravascular tissue is concentration-driven. Additionally, oxygen can diffuse from the arterioles to the venules. The amount of oxygen can be translated into oxygen saturation (*S*_*i*_*O*_2_), from which the level of oxygenated (*HbO*_*i*_) and deoxygenated blood (*HbR*_*i*_)can be estimated.

## Discussion

This work investigates the different contributions of inhibitory and excitatory neurons to the neurovascular response using experimental data from several different studies in a model-driven meta-analysis. This approach allows us to generate quantitative estimates for the cell-type-specific contributions to the fMRI-BOLD-signal. Our analysis shows that the inhibitory neurons are the biggest contributors to the BOLD-response (Figures 2A and D) in 8 of the 11 experiments (Vazquez, Fukuda and Kim, 2018; Lee *et al*., 2020; Moon *et al*., 2021), and always the biggest contributor to the vascular response (Figure 2C). Furthermore, the mode-based analysis accurately describes the key qualitative dynamics observed in data (Figures 3, 6, and S3), and correctly predict new validation data not used for model training: the response to NOS-inhibition (Figure 3G), and a 2 ms and 10 ms pulse width stimulation paradigm (Figure 4). Importantly, the model could describe how the inhibitory neurons act as the primary contributors to the regulation of the vascular response (Figure 3). Furthermore, the model was able to explain why high-intensity, but not low-intensity, stimulations of interneurons lead to a biphasic BOLD-response (Figure 6): it is because the stimulation of NO-interneurons (Figure 6G) is balanced out by GABA (Figure 6D), while strongly inhibiting the pyramidal cells, causing a net vasodilation (Figure 6J, black) as the release of NO (Figure 6K) cannot offset the lack of PGE_2_ (Figure 6L). Finally, we could use the model to explain why there is a double-peak in response to prolonged SOM stimulations (Figure 7A, crimson): a) the initial peak is generated by the direct vasoactive influence of the SOM-releasing neurons; b) the second peak is generated by a delayed recovery from the inhibition of activity in the other neuron populations (NO, NPY, and Pyramidal neurons)(Supplementary Materials section 2). In summary, our results provide a detailed, quantitative, and mechanistic underpinning for a new consensus view across 11 different experiments: that it is interneurons and not pyramidal cells that dominate the BOLD response.

### Other models investigating the neurovascular coupling

Various studies have developed different types of mathematical models to investigate the NVC (Griffeth and Buxton, 2011; Havlicek *et al*., 2015; Kim and Ress, 2016; Lundengård *et al*., 2016; Di Volo *et al*., 2019; Sten *et al*., 2020, 2023; Moon *et al*., 2021; Tesler, Linne and Destexhe, 2023). However, very few studies have investigated the more prominent role of inhibitory neuron populations suggested by recent experimental results. Firstly, the work by Moon *et al*. 2021 describe a 2-pole, 1-zero Laplace function, which primarily focuses on capturing the qualitative temporal features of the so-called hemodynamic response function (HRF)(Moon *et al*., 2021). However, this model offers no mechanistic insights into the neurovascular response. Secondly, Tesler *et al*. proposes a model that describes the contribution of astrocyte calcium dynamics to the relationship between neuronal activity and the fMRI-BOLD response (Tesler, Linne and Destexhe, 2023). This work uses a mean-field neuron spiking model (Di Volo *et al*., 2019) to describe selective activation of excitatory and inhibitory neurons. The Tesler model is limited by only having the astrocytes, thus not including the prominent effect of NO, controlling CBF and primarily focusing on excitatory neuronal stimulation. Lastly, our own prior work (Sten *et al*., 2023) presents a model including both inhibitory and excitatory neurons that could describe neurovascular control in response to different stimulation types, including optogenetic stimulations. Similarly to the other mentioned models, the Sten model predominantly focuses on excitatory neuronal stimulation as a main contributor to the regulation of CBF. As such, to the best of our knowledge, no studies are using state-of-the-art mechanistic models to investigate the qualitative differences (Figures 3-7, S3) of inhibitory neuronal populations to NVC, and no previous analysis has been able to quantify the cell-specific quantitative contributions to the BOLD, CMRO2, HBR, and CBV responses (Figures 2, S1-S2) to the NVC.

### Model explanation of key qualitative behaviours in data

Our model analysis, based on our mechanistic mathematical model (Figure 8), provides a cohesive framework to investigate the different qualitative behaviours seen in the experimental data. We present such explanations for: 1) the dominant neurovascular control of inhibitory neurons (Figure 3); 2) the biphasic response observed for high-intensity inhibitory stimulation (Figure 6); 3) the emergence of a double-peak behaviour stimulation SOM-interneurons (Figure S3).

To the first point, the dominant control of inhibitory neurons, our model captures this behaviour well over different data (Figure 3A-E). The simulations capture the prominent features in data: a peak value occurring roughly 9 seconds after the stimulation has ceased (0-4 sec), followed by a return to baseline. While explaining the change in HbO, HbR, HbT, and OIS-BOLD well (Figures 3A–C, E, orange), the model slightly overshoots the CBF response (Figure 3D, orange). Similarly, the model captures the more modest responses generated from the excitatory optogenetic stimulation (Figures 3A–E, blue). The model also captures the qualitative difference in HbR-dynamics between the different stimulations, i.e. decreasing during inhibitory stimulation and increasing during excitatory stimulation. The increase in HbR is attributed to a combination of a high metabolic load and a relatively weak vascular response (Figure 3D, blue). Model analysis reveals that the higher metabolic load is attributed to the stimulation of excitatory neurons, which have a greater contribution to CMRO_2_ (Figures 2B, S1)(Meyer *et al*., 2011) These different HbR-responses are also mirrored in the OIS-BOLD response (Figure 3E). Additionally, for the sensory stimulation (Figure 3, inserts), in which both the inhibitory and excitatory pathways are activated, we see again that the inhibitory neurons contribute the main part of the vascular response (Figures S1-S2, Exp. 2 and 3).

Secondly, our analysis of the stimulation-intensity-dependent biphasic behaviour (Figure 6), also suggests a distinct temporal hierarchy in neuronal activation. We observe a very rapid response from the NO-interneurons (Figure 6I and J, red); followed by a slightly delayed activation of the pyramidal neurons, which release PGE_2_ (Figure 6I-J, blue); and lastly by the considerably slower response NPY-interneurons. This dynamic of neuronal activation aligns well with the HRF-model presented by Moon *et al*., which describes both the fMRI-BOLD and HbR responses with a fast-acting inhibitory component and a slower-acting excitatory component (Moon *et al*., 2021). Additionally, the same temporal hierarchy of neuronal activation was also observed in our previous work (Sten *et al*., 2023), where this dynamic could be preserved across multiple experiments in different species. This alignment across different research efforts underscores the foundational nature of this temporal hierarchy. Moreover, our model extends these insights by offering a more detailed mechanistic explanation of the processes involved. Additionally, the stimulation-intensity-dependent model explanation also aligns with recent experimental findings presented by Dadarlat *et al*., where they also observe that the level of stimulation intensity affects the response of the inhibitory neurons (Dadarlat, Sun and Stryker, 2024).

Thirdly, the emergence of a double-peak behaviour, seen in Figure 7, is associated with the stimulation of SOM-interneurons (Lee *et al*., 2020). The double-peak dynamic could be explained by the SOM-interneurons having slower intracellular pathways, compared to NO-interneurons (Figure S3, Supplementary Materials section 2). Moreover, the model simulation suggests that the double-peak dynamic is attributed to a reduction and recovery of the vasoconstrictor NPY. In our previous work (Sten *et al*., 2023), we put forward another mechanism for describing the double-peak behaviour presented by Drew *et al*. (Drew, Shih and Kleinfeld, 2011). However, this previous explanation relied on the excitatory neurons acting as the primary driver of the neurovascular response. The data analysed in this work (Vazquez, Fukuda and Kim, 2018; Lee *et al*., 2020; Moon *et al*., 2021) do not support this excitatory-driven explanation. Instead, the current work puts forth an explanation where the inhibitory neurons are the main contributor to the neurovascular response.

### Model analysis limitations and critical features in data

The model analysis is limited to the information contained in the training data. The analysed optogenetic data are gathered from a range of varying experimental protocol parameters, see Supplementary Table 1, where stimulation strength, duration, frequency, and anaesthesia dosing protocol differ. Our framework attempts to analyse these data collectively to investigate how these differences relate to the experiments. This means that the analysis is subject to some limiting features presented in the experimental protocol and contradictory behaviour in the resulting data.

One example of contradictory behaviour are the Hb responses to stimulating the NO-interneurons in Lee *et al*. (Figure 7, light red)(Lee *et al*., 2020), compared with the high-intensity stimulation of NO-interneurons (Figure 5, yellow)(Moon *et al*., 2021). These experiments share similar stimulation protocol parameters (Supplementary Table 1) and therefore one would expect similar vascular responses. However, the Hb dynamics differ greatly and most notable is that the biphasic response is missing from the Lee *et al*. NO-response. Instead, the NO-response is qualitatively closer to the 1 Hz stimulation response presented by Moon *et al*. (Figure 5, orange). It should be noted that these studies use different optogenetic lines (NOS-connected pathway in Lee *et al*. and vesicular GABA transport (VGAT) vectors in Moon *et al*.) to introduce the light-sensitive Channelrhodopsin-2 (ChR2) ion channels into the mice. While this choice of vector could be expected to cause some differences in the vascular responses, one would not expect such qualitatively different responses. Rather, these differences in responses could be attributed to the data being recorded in different brain areas and/or different anaesthesia protocols being used.

Factors that affect the stimulation and how the stimulation is perceived by the cells, e.g. anaesthesia protocol, could contribute to the qualitative different response seen across the three analysed studies (Vazquez, Fukuda and Kim, 2018; Lee *et al*., 2020; Moon *et al*., 2021). For example, when observing the overall light exposure per second for each study, analysis reveal that all optogenetic experiments employ 150-200 ms of light exposure per second (Supplementary Table 1). However, these similar light exposures cause widely different neurovascular responses (Figures 3-7) and the differences probably lie in how the electrophysiological activity responds to the different stimulation protocols. For instance, to explain the frequency-dependent response seen in Figure 6 we needed a stimulation saturation effect. Such a mechanism could be connected to the recovery phase present in electrophysiological models, i.e. the hyperpolarization phase of neuronal ion channels (Tsodyks and Markram, 1997; Rosenbaum, Rubin and Doiron, 2012) and global depressive phases of neuronal networks (Zhang *et al*., 2016). These systems actively regulate themselves to avoid overstimulation and our model analyses indicate a similar role for GABA (Figure 6D). Furthermore, it has been shown that the anaesthesia itself influences NVC dynamics (Uhlirova *et al*., 2016). We have previously presented a modelling approach to investigate the anaesthesia effect on the vascular control (Sten *et al*., 2020). However, to the best of our knowledge, no such investigation has yet been performed on the electrophysiological properties in response to the optogenetic stimulation. If one were to analyse the interaction of anaesthesia and optogenetic stimulation in *in silico* spiking neuronal network models (Lee, Wang and Hudetz, 2020; H. Lee *et al*., 2021), one could potentially find explanations for the observe vascular differences.

We should note that our model structure does not attribute any vascular control to the astrocytes, instead they only contribute to the metabolism. As the model was iteratively developed (Figure 1D), the astrocytes were initially included solely to regulate the levels of glutamate and GABA, and subsequently the activity of the neurons. However, the model also required the metabolic contributions of the astrocytes to accurately explain the data, while their direct vascular regulation could still be assumed to be negligible. This does not necessarily mean that the astrocytes’ contributions to vascular regulation are negligible, only that the current model structure could explain the considered data given this assumption. This assumption stands in contrast to other modelling frameworks, such as Tesler *et al*. (Tesler, Linne and Destexhe, 2023), that do include astrocytes with an active role in vascular control. Similar assumptions have also been made for other interactions between the astrocytes and neurons.

Lastly, we acknowledge limitations in the level of detail of our neural model that could be addressed to improve our predictions. For example, the model only includes one type of excitatory (pyramidal) and three types of inhibitory neurons (SOM, NPY, NO), but other known neuron types and subtypes such as interneurons expressing parvalbumin (PV) or vasoactive intestinal peptide (VIP), may play important roles in NVC (Krogsgaard *et al*., 2023; Yao *et al*., 2023). Additionally, the modelled spatial distribution and synaptic connectivity of neurons is currently overly simplified, and omits, for example, the six layer cortical structure and complex interlaminar and long-range connectivity patterns (Harris and Shepherd, 2015). These spatial and connectivity properties may have important implications for calculating NVC responses, as local neural activity may trigger strong responses in distally connected neural populations. Polysynaptic inhibitory connectivity patterns, such as disynaptic inhibition (E->I->E) and disinhibition (I->I->E), may be important to interpret the contribution of inhibitory activity in the BOLD response. We are working towards extending our analysis using data-driven multiscale biophysical circuit models that incorporate known cell types, anatomical structure and synaptic connectivity (Dura-Bernal *et al*., 2019; Dura-Bernal, Griffith, *et al*., 2023; Dura-Bernal, Neymotin, *et al*., 2023).

### An emerging consensus view that inhibitory neurons dominate the vascular and BOLD response: arguments for and against

The result of this work primarily argues for a new consensus view that inhibitory neurons, and not pyramidal neurons, are the most dominant cell type in generating vascular and BOLD responses. We have quantified the contribution of the different cell types and found that interneurons dominate the BOLD response in 8 out of 11 experiments, and the vascular response in all 11 experiments (Figure 2). Furthermore, we have provided an extensive mechanistic underpinning for how this new role of interneurons can be used to explain various details in responses, such as a biphasic response to high-intensity interneuron stimulation (Figure 6), and a double-peak response to prolonged SOM stimulation (Figure 7). All of these results argue for a new and more prominent role of interneurons, which is compatible with many new studies (Vazquez, Fukuda and Kim, 2018; Echagarruga *et al*., 2020; Moon *et al*., 2021). However, it should also be noted that other, and contradictory results, also have been reported. For instance, studies on excitatory neuron stimulation show both a modest (Vazquez *et al*., 2014; Vazquez, Fukuda and Kim, 2018) versus prominent (Desjardins *et al*., 2019; J. Lee *et al*., 2021; Moon *et al*., 2021) hemodynamic responses. It has also been shown that optogenetic excitatory stimulation can yield a stronger neurovascular response compared to optogenetic inhibitory stimulation (Desjardins *et al*., 2019). In summary, it is thus clear that our work provides a step towards forming a new consensus view, but that further work needs to be done to get a view that truly encompasses all the seemingly contradictory data.

## Method

To analyse the different dynamics of the NVC that are not unified with the conventional interpretation of vasoactive regulation, a mechanistic mathematical model has been used. This type of model allows for greater integration of experimental data from multiple different studies, considering multiple different variables such as OIS-BOLD, fMRI-BOLD, and Hb levels. The model developed in this study consists of ordinary differential equations (ODEs) and differential algebraic equations (DAEs) and has been formulated to describe the mechanisms involved in the NVC. Parameter estimation, identifiability, and model observability analyses were performed, and the model has been evaluated with respect to its ability to accurately describe different sets of experimental data. These analyses are described in detail in this section.

### Model formulation

The model used in this work is formulated as a system of ODEs and DAEs and can in general be formulated as:

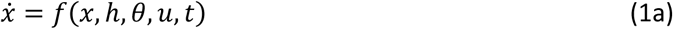

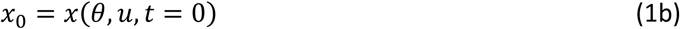

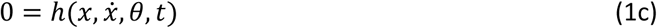

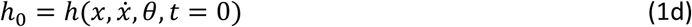

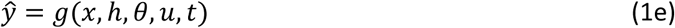

where *x* is a vector containing the model states that can be derived with respect to time *t*; 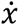 are the derivatives of states *x* with respect to *t*; *x*_0_ is the values of the states *x* at the initial timepoint *t* = 0; *θ* is a vector with the model parameters; *u* specifies the model inputs; *f*, *h*, *g* are non-linear smooth functions; *f* describes how the derivatives of the states *x* change over time with respect to the current value of *x*, the parameters *θ*, and the input *u*; *h* describes the system of differential equations and algebraic equations that have no corresponding time derivatives; *h*_0_ are the initial values of the algebraic expressions at time point *t* = 0; and *ŷ* is a vector of the model’s observable properties which is described by the function *g*.

### Model structure

The model used in this study is derived from the model presented in our previous work (Sten *et al*., 2023). This section will give a brief description of this model structure and a detailed description of what changes have been made to the model structure for the work presented herein. For a detailed description of the original model please see the original publication (Sten *et al*., 2023) and the Supplementary Materials section 5.

### Expansion of the Sten *et al*. 2023 model

The original model presented in Sten *et al*. 2023 was expanded to accurately explain the experimental data considered herein. These expansions include but were not limited to, an addition of a readout for the OIS-BOLD signal, an expanded expression for the oxygen permeability of the blood vessels and tissue uptake of oxygen, and the introduction of saturated neuronal activity. These additions to the model are detailed in the sections below.

#### Addition of BOLD OIS signal

The OIS-BOLD signal was implemented in the model as an exponential expression of the relative change in HbR and CBV levels. The HbR component represents the approximation of the OIS-BOLD signal as an inverse measurement of the change in HbR. The volume-based offset was introduced to account for the light scattering effect caused by an increased blood volume, i.e. an increased blood volume leads to increased light scattering which means a decrease in optical signal. This relationship is explained by the following expression:

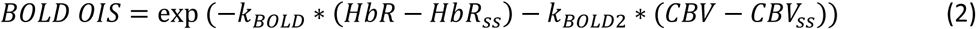

where *k*_*BOLD*_ and *k*_*BOLD*2_ are scaling parameters which are estimated from data; *HbR* is the calculated levels of deoxygenated haemoglobin; *CBV* is the calculated cerebral blood volume; and *HbR*_*ss*_ and *CBV*_*ss*_ are the baseline values of HbR and CBV, respectively.

#### Oxygen permeability and tissue oxygen uptake

The original model had a simplified explanation of the blood vessels’ oxygen permeability and tissue oxygen uptake. This expression needed to be expanded for the model to describe the data since the simulated values for the increases in CBF and relative change in HbO did not quite reflect the observed changes in the data (Vazquez, Fukuda and Kim, 2018). This expanded expression for the oxygen permeability was implemented as two separate mechanisms from the model presented by Barret *et al*. 2015 (Barrett and Suresh, 2015) the first mechanism represents a dynamic permeability of the capillary vessels, formulated as:

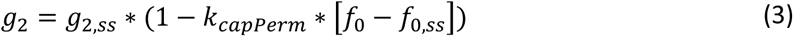

where *g*_2_ is the permeability of the capillary vessels; *g*_2,*ss*_ is the permeability at baseline; *k*_*capPerm*_ is a scaling parameter; *f*_0_ is the blood flow entering the local region; and *f*_0,*ss*_ is the blood flow at baseline.

The second mechanism describes the effect of recruiting upstream oxygen i.e., the mechanism introduces a “leakage” of oxygen from the blood to other tissue before reaching the brain. In the original model, this leakage is considered to be constant for the duration of the stimulation. In the updated model a dynamic leakage is implemented as:

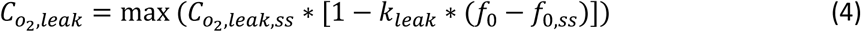

Where 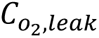 is the loss of O_2_ concentration due to “leakage”; 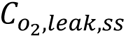 is the leakage at baseline; *k*_*leak*_ is a scaling parameter; and *f*_0_ and *f*_0,*ss*_ is the blood flow and basal blood flow as described above. The max(x, 0) function is to prevent a negative O_2_ leakage i.e., the blood entering the local region of the brain cannot have a higher O2 concentration than the systemic arteries.

#### 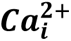 prevented from becoming negative

The original model allowed the states that represented the neuronal *Ca*^2+^levels to become numerically negative if the effect of the corresponding neuron activity was negative enough. Negative enough means that during stimulation the state of the neuronal activity *N*_*i*_ becomes so negative that its contribution overtakes the corresponding 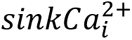-term and the 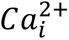-state becomes negative. This was addressed in the expanded model by introducing a softplus activation function in the equation for the 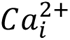-states, to prevent the inflow term that corresponds to stimulation from the neuronal activity *N*_*i*_ to be smaller than 0. Meaning that the inflow of *Ca*^2+^will never go below the baseline. This means that the differential equation for the 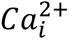 states are formulated as:

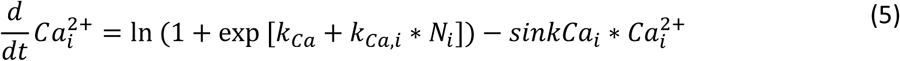

where 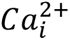 is the collective level of *Ca*^2+^for neuron type *i*; *k*_*Ca*_ and *k*_*Ca*,*i*_ are general and neuron-type-specific scaling parameter respectively; *N*_*i*_ is the neuronal activity of neuron type *i*; and 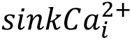 is a collective term for the reduction in the *Ca*^2+^levels.

#### Changed CMRO_2_ dependencies

In the original model, the calculation of the overall CMRO_2_ depended on the activity of the different neuron populations *N*_*i*_. However, much like the 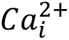states, this becomes a problem if *N*_*i*_ becomes negative. Therefore, the expression for CMRO_2_ was changed to be proportional to 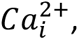 rather than *N*_*i*_. Additionally, individual scaling parameters for the different neuron populations contributions to the total CMRO_2_ were introduced. This resulted in the following expression for CMRO_2_:

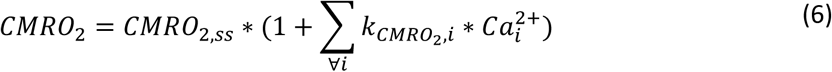

Where *CMRO*_2_ is the local region’s overall metabolic rate of O_2_; *CMRO*_2,*ss*_ is the metabolic rate at baseline; 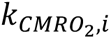 is the scaling parameter for the metabolic contribution of neuron population *i*; and 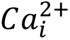 are the *Ca*^2+^ levels for neuron population *i*.

#### Reformulated intraneuronal signalling, introduced GABA and Glutamate State and a saturated neuronal activity ***N*_*i*_**

In the original model, the effects of neurotransmitters glutamate and GABA (i.e., inhibitory/excitatory pathways) are integrated in the states that represent the neuronal activity *N*_*i*_. To make the model more readable and to better account for the excitatory and inhibitory regulation between, and within, the neuronal populations the expanded model introduces additional states for glutamate and GABA. The glutamate state increases the different neuronal activities *N*_*i*_, while the GABA state decreases neuronal activities (Figure 8A). Additionally, a saturating effect was added to the neuronal activity induced by the experimental stimulations, meaning that for prolonged high-intensity stimulations, the increase in neuronal activity caused by the stimulation is capped to the value determined by the parameter estimation. All in all, these reformulations, and additions to the neuronal signalling mean that for the expanded model the equations for *N*_*i*_ states are formulated as:

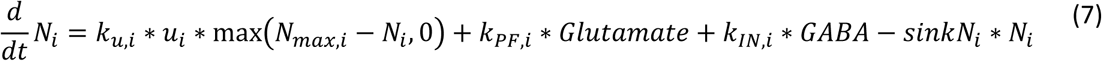

where *u*_*i*_ is the stimulation function; *N*_*max*,*i*_ is the parameter determining the saturation in the stimulating effect i.e., the stimulation term becomes 0 when *N*_*i*_ > *N*_*max*,*i*_; *glutamate* and *GABA* are the changes in glutamate and GABA levels respectively; *k*_*u*,*i*_,*k*_*PF*,*i*_, *k*_*IN*,*i*_, and *sinkN*_*i*_ are scaling parameters scaling the effects of the stimulation, the neuronal activation caused by glutamate, the neuronal inhibition caused by GABA, and the general return to baseline neuronal activity, respectively. The levels of glutamate and GABA are implemented as

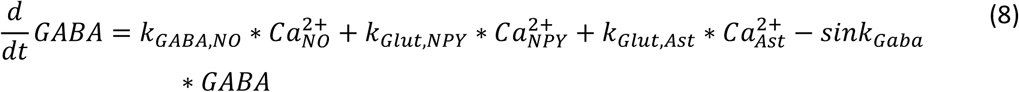

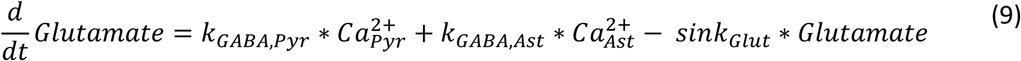

Where *k*_*Glut*,*i*_ and *k*_*GABA*,*i*_ are kinetic rate parameters determining the release of glutamate and GABA from the respective neuron populations; 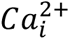 are the *Ca*^2+^ levels for the respective neuron populations; *sink*_*i*_ is a parameter determining the rate at which the glutamate and GABA levels return to baseline.

The stimulation function *u*_*i*_ is implemented as a repeating step function that alternates between 0 and 1 depending on the stimulation scheme.

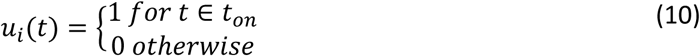

where *t*_*on*_ is the set of time points determined by the stimulation pulse frequency, pulse width, and total stimulation time. For example, a 4 second simulation at 1 Hz with a pulse width of 10 ms, *t*_*on*_will include the 10 first milliseconds of each second of the stimulation time.

#### Addition of astrocytes as a regulating force for Glutamate and GABA levels

In addition to the reformulation of the glutamate and GABA dynamics, the expanded model also has a state representing the activity in astrocytes/glial cells. This state was introduced to allow the model a higher degree of flexibility in regulating the levels of glutamate and GABA. The astrocyte activity is formulated in the same way as equation 7. Although since there is no external stimulation of the astrocytes the corresponding *k*_*u*,*Ast*_ = 0 i.e., the term representing the stimulation is removed.

### Model evaluation

The model was evaluated with respect to its ability to describe experimental data taken from published literature. This evaluation consisted of estimating the model parameters through iterative fitting to part of the data, evaluating the model fit with statistical testing, analysing parameter identifiability and model observability, and lastly validating model predictions against independent validation data not used for model training.

### Parameter estimation

The model’s parameters were estimated by iteratively fitting the model to the training data. This was achieved by using an optimization algorithm to minimize the sum of the squared residuals between the model simulation and the experimental data. The squared residuals were, where applicable, weighted by the Standard error of the mean (SEM) of the data. Thus, the function that was minimized was formulated as:

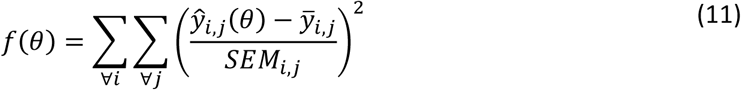

were 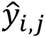 is the model simulated value of variable *i* at timepoint *j*; *θ* are the model parameters; 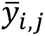 is the mean data point of variable *i* at timepoint *j*; and *SEM*_*i*,*j*_ is the standard error of the mean for variable *i* at timepoint *j*. The function *f*(*θ*) was implemented as the objective function for a general optimisation algorithm and minimised with respect to the parameters *θ*, such that:

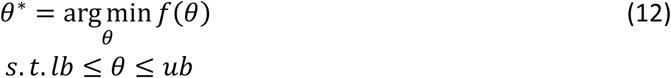

where *θ*^∗^are the optimal parameter values i.e., the parameter values that yield the best agreement between the model and the data; and *lb* and *ub* are the lower and upper bounds of the parameter estimation search space respectively.

### Identifiability analysis

The identifiability of the model parameters was determined by applying a Markov Chain Monte Carlo (MCMC) sampling. An adaptive multi-chain implementation of the Metropolis-Hastings algorithm, using 10^6^ samples, were used to generate a posterior distribution of parameter values. All parameter sets *θ* that were deemed acceptable with respect to a *χ*^2^-test was saved and the practical parameter identifiably for each parameter *θ*_*i*_ were determined by the confidence interval:

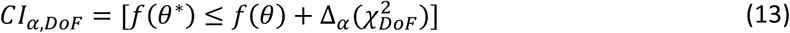

were *α* is the significance level of the *χ*^2^-test; *DoF* is the number of degrees of freedom for the model; *f*(*θ*^∗^) is the solution for the objective function in eq. 2 for the optimal parameters *θ*^∗^; Δ_*α*_ is the *α* quantile of the 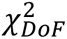 statistic (Kreutz *et al*., 2013; Maiwald *et al*., 2016). For the analysis conducted in this study, the *DoF* was set to the number of data points subtracted by the number of model parameters. The estimated values for the optimal parameter values and the parameter confidence intervals can be found in Supplementary Tables 2 and 3.

### Model prediction of smaller pulse widths

In Figure 4, the model predictions of the 2 ms and 10 ms pulse width stimulation protocol, presented by Vazquez *et al*. 2018 (Vazquez, Fukuda and Kim, 2018), are depicted. As the stimulation does not linearly affect the system, or model, we allowed the parameters governing the stimulation effect, *k*_*u*1_, *k*_*u*2_, and *k*_*u*3_, to be scaled for these predictions. Note that no other parameters were altered, and the scaling does not amount to re-training of the model. Rather, the scaling is considered a mechanism for interpreting the effects of the stimulation. The scaling factor is described below.

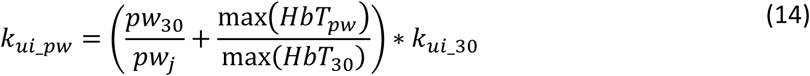

Where, *k*_*ui*_*pw*_ are the rescaled parameters for the different pulse widths (*pw*), *i* = {1, 2, 3} and *pw* = {2, 10}, *HbT*_*pw*_ are the mean values of the HbT data for the different pulse widths, *HbT*_30_ is the mean value for the HbT data corresponding to the 30 ms experiment, and *k*_*ui*_30_ are the estimated model parameters, obtained from the model training.

### Experimental data

The main computational analysis of the work presented here considered experimental data from three previously published studies. The first of which was presented by Vazquez *et al*. in 2018, where they investigated the contribution of different types of neurons to the hemodynamic response and the metabolic load of the NVC (Vazquez, Fukuda and Kim, 2018). The second study, presented by Moon *et al*. in 2021, further investigated the contributions of excitatory and inhibitory types of neurons to the fMRI-BOLD response (Moon *et al*., 2021). The third and final study was presented by Lee *et al*. in 2020 and investigated the hemodynamic response when selectively stimulating different subpopulations of inhibitory interneurons (Lee *et al*., 2020). All three of these studies used optogenetic mice to selectively stimulate different types of neurons. That is the use of mice that have been genetically engineered to express the light-sensitive Channelrhodopsin-2 (ChR2) ion channel in specific neuron populations (Nagel *et al*., 2003; Arenkiel *et al*., 2007). The light-sensitive properties of ChR2 mean a light pulse can be used to open the ion channel and thus selectively activate specific neuron populations (Fenno, Yizhar and Deisseroth, 2011). All the data used in this work were sourced from figures published in previous studies.

### Data describing the neurovascular response for excitatory and inhibitory stimulation

In their study from 2018, Vazquez *et al*. used OIS imaging to measure the relative levels of oxygenated Hb (HbO), reduced or deoxygenated Hb (HbR), and the total amount of Hb (HbT) in the somatosensory cortex of anesthetized mice. They also use laser doppler flowmetry (LDF) to measure CBF. In this study, they consider data from 41 transgenic mice, 22 of which had inhibitory neurons expressing ChR2 via the vesicular GABA transporter (VGAT)(Zhao and Zhang, 2011; Vazquez, Fukuda and Kim, 2018), and 19 had excitatory neurons expressing ChR2 via the thymus cell antigen 1 (Thy1) promoter (Vazquez *et al*., 2014). The mice were exposed to optogenetic stimulation consisting of light pulses with an amplitude of 1 mW, a frequency of 5 Hz, and pulse widths of 2 ms, 10 ms, and 30 ms were evaluated. The total stimulation time for the optogenetic stimulation was 4 seconds. The authors also evaluated the hemodynamic response to sensory forelimb (FL) stimulation by applying a 1.2 mA, 4Hz electric pulse for 0.5 ms to one of the mice’s forepaws. The mice were anesthetized using an initial dose of ketamine (75 mg/kg) and xylazine (10 mg/kg), and a continuous supply of ketamine (30 mg/kg/h) during the experiments. For further details please see the original publication (Vazquez, Fukuda and Kim, 2018). The data considered in this present work mainly comes from the experiment entitled “*Experiment 5*” in the original publication that investigates the differences in the hemodynamic responses and CMRO_2_ invoked by optogenetic photo-stimulation of excitatory and inhibitory neurons.

### Data illustrating the effects of stimulation intensity on the neurovascular response

Building on the findings published by Vazquez *et al*. in 2018 (Vazquez, Fukuda and Kim, 2018), in their study from 2021 Moon *et al*. investigate the contributions of excitatory and inhibitory neuronal activity to the fMRI-BOLD response (Moon *et al*., 2021). In this study, the authors measured the fMRI-BOLD response for both excitatory, inhibitory, and sensory stimulation from the different regions of the brain in optogenetic mice. OIS imaging was also used to quantify changes in the levels of HbO, HbR, and HbT for inhibitory stimulation. This study used 17 VGAT-ChR2 mice for inhibitory stimulation, 6 mice that expressed ChR2 via the calcium-calmodulin-dependent protein Kinase II (CaMKII) gene for excitatory stimulation, and 5 wild-type non-transgenic C57BL/6 mice as a control group. For the fMRI-BOLD measurement, the mice were ontogenetically stimulated with 200 ms and 10 ms light pulses at frequencies of 1 Hz and 20 Hz respectively, for a total stimulation length of 20 s. For the sensory stimulation, an electrical current of 0.5 mA at 4 Hz was applied to the left forelimb for 20 seconds. The fMRI measurements were collected at a magnetic field strength of 15.2T, with a repetition time (TR) of 310 ms, an echo time (TE) of 3 ms, and a flip angle (FA) of 30°. For the OIS measurements the same optogenetic stimulation as for the fMRI experiment were used i.e., 200 ms and 10 ms light pulses at 1 Hz and 20 Hz, respectively. For the OIS measurements two different stimulation times were used, namely one longer 20 s stimulation and one shorter 5 s stimulation. In this study the mice were anesthetized with an initial dose of ketamine (100 mg/kg) and xylazine (10 mg/kg), and a continuous supply of 25 mg/kg ketamine and 1.25 mg/kg xylazine during the experiments. For further details please see the original publication (Moon *et al*., 2021).

### Data describing contributions to neurovascular response from different inhibitory interneuron populations

In their study published in 2020, Lee *et al*. investigate the differences in the NVC’s hemodynamic response if different subpopulations of GABAergic interneurons are stimulated. In this study, the hemodynamic responses in mice for stimulating NO-interneurons and SOM-interneurons were compared with each other, and with a sensory whisker stimulation. OIS imaging where used to measure the relative changes in the levels of HbT, HbO, and HbR. The study cohort consisted of a total of 39 mice, male and female. 18 of these mice were nNOS-CreER × ChR2-EYFP (nNOS-ChR2) mice, selectively expressing ChR2 in NO-interneurons. 12 mice were Sstm2.1Crezjh/j × ChR2-EYFP (SST-ChR2) mice, selectively expressing ChR2 in NO-interneurons and SST interneurons. Lastly, 5 non-transgenic C57BL/6J mice and 4 non-ChR2 expressing nNOS-ChR2 mice were used for sensory stimulation. The optogenetic stimulation paradigms consisted of 10 ms light pulses at 20 Hz applied to the subjects for shorter 2 s and longer 16 s stimulation periods, respectively. For the sensory stimulations, the whiskers were deflected at 5 Hz for either 2 or 16 s. The animals were anesthetized with an initial injection of fentanyl-fluanisone and midazolam diluted in water (7 ml/kg), and a continuous supply of isoflurane in oxygen at 0.8 L/min during the experiments. Please see the original publication for further details (Lee *et al*., 2020).

### Model Implementation

The mathematical model described in the “Model structure” section above was implemented and simulated using the MATLAB version of the “Advanced Multi-language Interface to CVODEs and IDAs” (AMICI) toolbox (Fröhlich *et al*., 2021). The parameter estimation and model evaluation were done using the “Metaheuristics for Systems Biology and Bioinformatics Global Optimization” (MEIGO) toolbox for MATLAB (Egea *et al*., 2014). Within the MEIGO framework, the enhanced scatter search (eSS) global algorithm and the dynamic hill climbing (dhc) local algorithm were used to solve the optimization problem formulated in the section “Model Evaluation”. For the identifiability analysis, the MATLAB implementation of the “Parameter EStimation TOolbox” (PESTO) (Stapor *et al*., 2018) was used for the MCMC sampling. This implementation used a parallel tempering algorithm (Ballnus *et al*., 2018) to generate the posterior parameter distributions. The area under the curve calculations presented here were calculated using a numerical integration via a trapezoidal method. All the computational implementations of the model and analyses described above were done in MATLAB by MathWorks Inc. releases R2017b and R2021b. All model implementation and files required to reproduce the work presented herein can be found at: https://github.com/Podde1/MetaAnalysisNVCInhibitoryNeurons.git and a permanent copy available at Zenodo (DOI: <u>10.5281/zenodo.13747286</u>).

## Acknowledgements

The computational resources required for these analyses were provided by the National Supercomputer Centre (NSC), funded by Linköping University.

## Author Contributions

**Conceptualization:** NS, HP, SS, ME, SDB, and GC. **Formal model analysis:** NS and HP. **Data curation:** NS and HP. **Visualization:** NS and HP. **Supervision:** GC, SS, ME, and SDB. **Funding acquisition:** GC, ME, and SDB.

All authors were involved in writing and reviewing the manuscript. All authors read and approved the final manuscript.

## Abbreviations

AUC: area under the curve
BOLD: blood oxygen level-dependent
CBF: cerebral blood flow
CI: confidence interval
CMRO_2_: cerebral metabolic rate of oxygen
fMRI: functional magnetic resonance imaging
GABA: *γ*-aminobutyric acid
Hb: haemoglobin
HbO: oxygenated haemoglobin
HbR: deoxygenated haemoglobin
HbT: total haemoglobin
HRF: hemodynamic response function
LDF: laser doppler flowmetry
NO: nitric oxide
NOS: nitric oxide synthase
NPY: neuropeptide Y
NVC: neurovascular coupling
OIS: optical intrinsic signal
PGE_2_: Prostaglandin E_2_
SOM: somatostatin
VGAT: vesicular GABA transport

## Supplementary Figures

**Figure S1:**
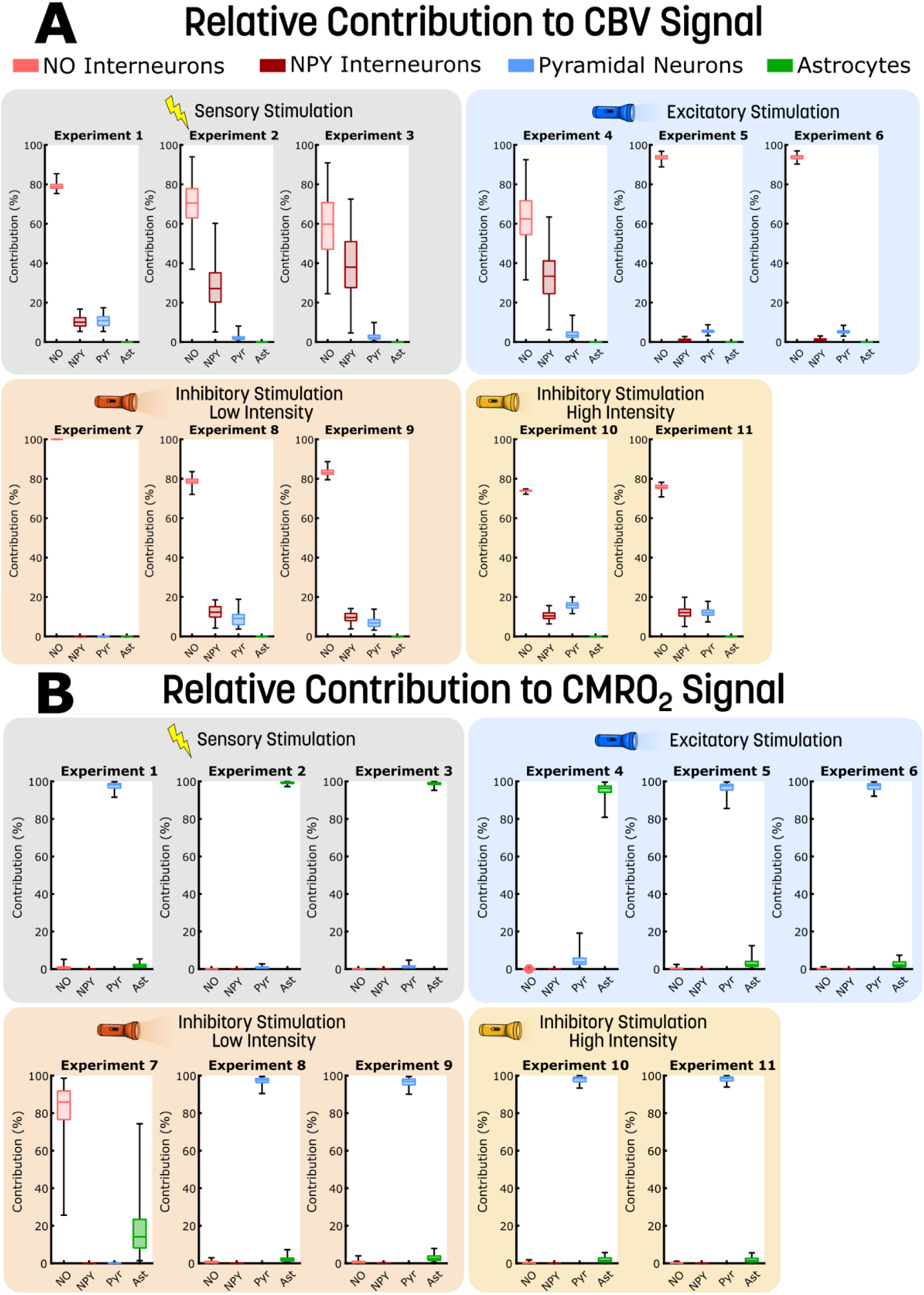
The contribution of different neuronal populations to the CBV and CMRO_2_ signal. The contribution of four different neuronal populations, nitric oxide (NO) interneurons (light red), neuropeptide Y (NPY) interneurons (crimson), pyramidal neurons (Pyr, blue), and glia cells (Ast, green), are presented for 11 different experiments to **A**) cerebral vascular volume (CBV) and **B)** cerebral metabolic rate of oxygen (CMRO_2_). The contributions are presented as boxplots, detailing the contribution of the four neuronal populations with uncertainty. The background of each experiment indicates the type of stimulation that generated these behaviours: grey is a sensory stimulation, blue is an optogenetic excitatory stimulation, orange is an optogenetic inhibitory low-intensity stimulation, and yellow is an optogenetic inhibitory high-intensity stimulation.

**Figure S2:**
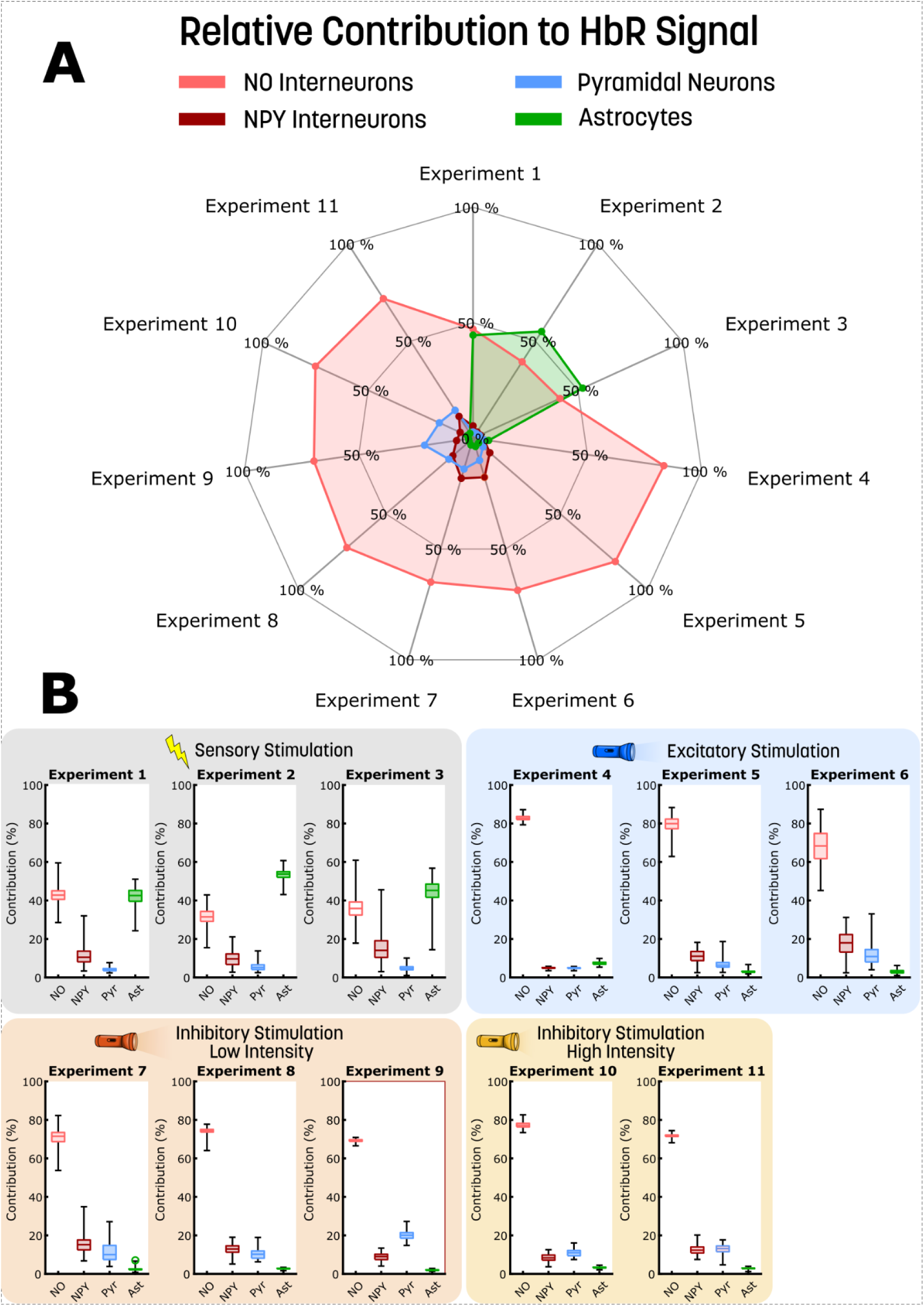
The contribution of different neuronal populations to the HbR signal. **A)** The contribution of four different neuronal populations, nitric oxide (NO) interneurons (beige), neuropeptide Y (NPY) interneurons (red), pyramidal neurons (Pyr, light blue), and glia cells (Ast, green), to the deoxygenated haemoglobin (HbR) are presented for 11 different experiments. **B)** Boxplots detailing the contribution of the four neuronal populations (NO beige, NPY red, Pyr light blue, Ast green), with uncertainty, for the 11 experiments presented in A. The background of each experiment indicates the type of stimulation that generated these behaviours: grey is a sensory stimulation, blue is an optogenetic excitatory stimulation, orange is an optogenetic inhibitory low-intensity stimulation, and yellow is an optogenetic inhibitory high-intensity stimulation.

**Figure S3:**
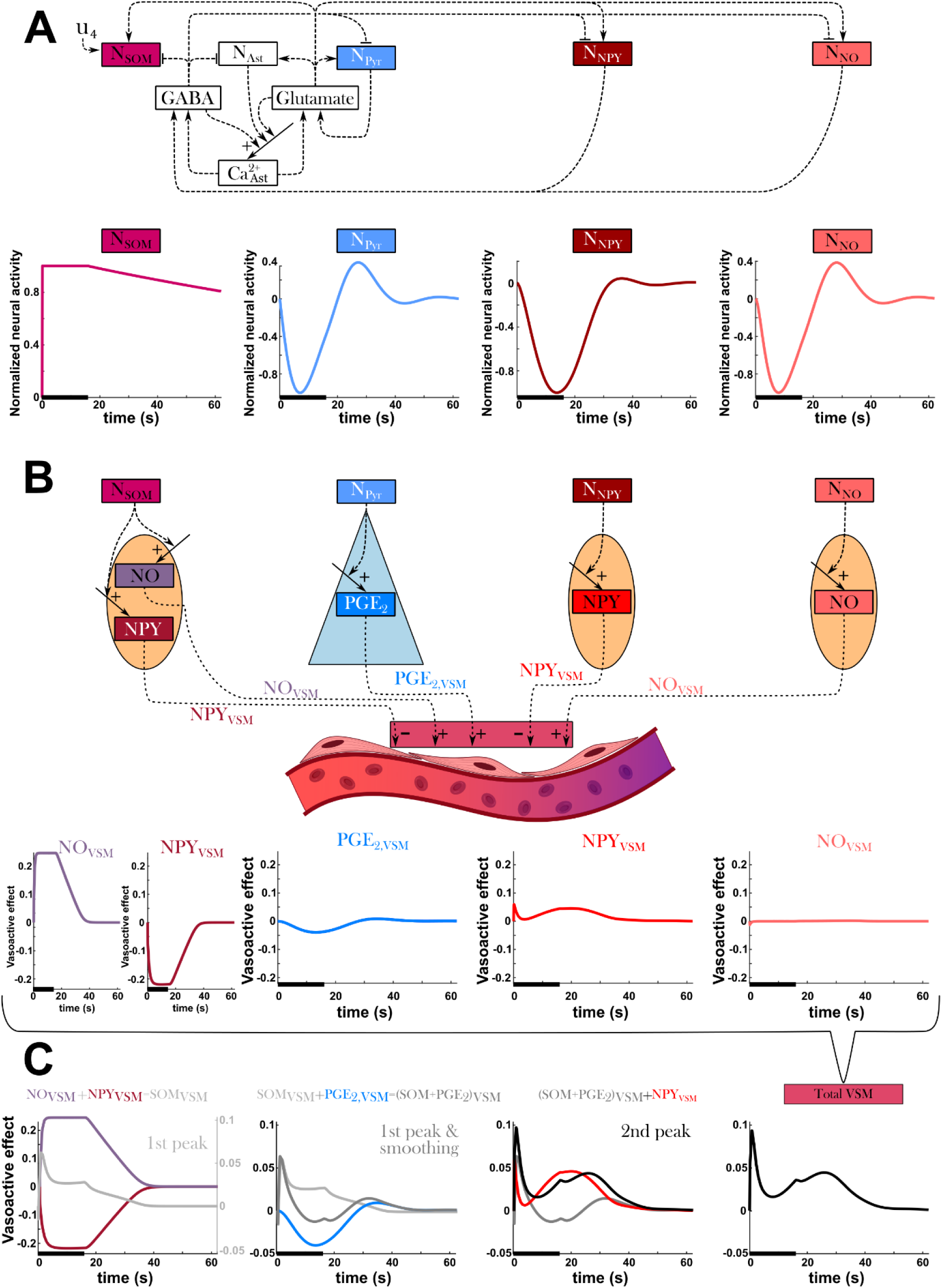
The delayed dilation of somatostatin-releasing interneurons is caused by different neuronal recovery speeds post-stimulation. **A)** A simplified overview of neural activity part of the model, including the extension of the somatostatin-releasing interneurons (SOM), for a 16 s long stimulation, u_4_, applied to the SOM neurons. As the SOM neurons (ruby) are stimulated, GABA is produced which inhibits the other neurons which are not externally stimulated. This is further highlighted by the plots of the normalized neural activity for SOM (ruby), pyramidal (blue), NPY expressing (crimson), and NO expressing (light red) neurons. For these plots, the x-axis represents time in seconds. **B)** A simplified model overview of how the neural activity propagates to the vasoactive signalling substances. NO and PGE2 dilate the vasculature, while NPY contracts. For the same 16 s long stimulation the vasoactive effect of NO (purple) and NPY (dark red), released by SOM neurons, PGE2 (blue), released by pyramidal neurons, NPY (red), released by NPY interneurons, and NO (light red), released by NO interneurons, are shown. **C)** By sequentially adding the individual vasoactive effects together, we can identify the contribution of the different substances. NO and NPY released by the SOM neurons produces the initial peak and a plateau that returns to the baseline (grey line). Adding PGE_2_ to the grey line transforms the plateau to the initial part of the second peak (dark grey line). Adding NPY released from the NPY interneurons further accentuates the initial peak and raises the amplitude of the second peak. NO released from the NO interneurons exhibit a weak impact on the total vasoactive response.

## Supplementary Table Legends

***Table S1: Overview of the experimental protocol parameters.*** The experimental stimulation protocol parameters are detailed for the 17 experiments analysed. The table includes the study of origin, stimulation type, stimulation duration, stimulation frequency, stimulation pulse duration in milliseconds and seconds, light exposure, stimulation strength or effect, the type of mouse and the utilized mouse strain, and the dosing of the anaesthesia agents. The colours indicate the stimulation type, where: grey is sensory stimulation, light blue is optogenetic excitatory stimulation, orange is optogenetic inhibitory low-intensity stimulation, yellow is optogenetic inhibitory high-intensity stimulation, and red is optogenetic somasomatic stimulation.

***Table S2: Model parameter values:*** The model parameters are detailed in this table in log10 scale. The columns detail: the parameter name, the best estimated value found during parameter estimation, the lower bound used during parameter estimation, the upper bound used during parameter estimation, the lowest value found for the confidence interval, and the highest value found for the confidence interval. Repeated parameters are named with an abbreviation to indicate which study and experiment they belong to.

***Table S3: SOM model parameter values:*** The model parameters are detailed in this table in log10 scale. The columns detail: the parameter name, the best estimated value found during parameter estimation, the lower bound used during parameter estimation, the upper bound used during parameter estimation, the lowest value found for the confidence interval, and the highest value found for the confidence interval. Repeated parameters are named with an abbreviation to indicate which experiment they belong to.

