## Supplementary Information for "A Model-Driven Meta-Analysis Supports the Emerging Consensus View that Inhibitory Neurons Dominate BOLD-fMRI Responses"

### 1 Cell-specific Relative Contribution to CBV, CMRO_2_, and HbR

Like the analysis of the cell-specific relative contribution to the BOLD signal, the relative contribution to the cerebral blood volume (CBV), the cerebral metabolic rate of oxygen (CMRO_2_), and the deoxygenated haemoglobin (HbR). These three variables are shown in the figures, Figures S1-S2, below.

In all the 11 experiments the vascular regulation, regulation of CBV, is dominated by the nitric oxide (NO) interneurons, regardless of the stimulation type (Figure S1A). This means that vascular changes of both the sensory stimulation type and the optogenetic excitatory stimulation type are attributed mainly to the NO interneurons.

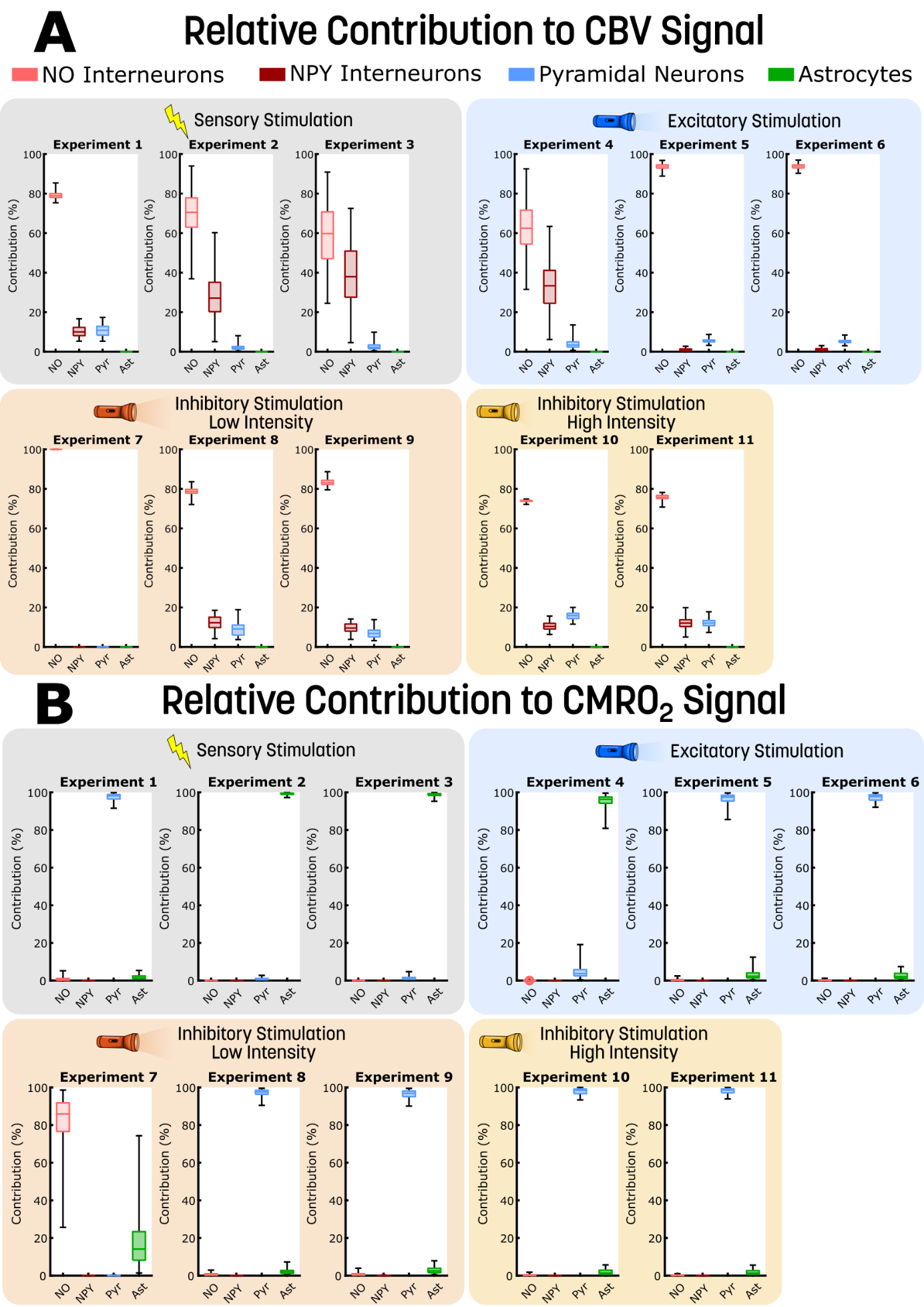

***Figure S1: The contribution of different neuronal populations to the CBV and CMRO_2_ signal.*** *The contribution of four different neuronal populations, nitric oxide (NO) interneurons (light red), neuropeptide Y (NPY) interneurons (crimson), pyramidal neurons (Pyr, blue), and glia cells (Ast, green), are presented for 11 different experiments to* ***A****) cerebral vascular volume (CBV) and* ***B)*** *cerebral metabolic rate of oxygen (CMRO_2_). The contributions are presented as boxplots, detailing the contribution of the four neuronal populations with uncertainty. The background of each experiment indicates the type of stimulation that generated these behaviours: grey is a sensory stimulation, blue is an optogenetic excitatory stimulation, orange is an optogenetic inhibitory low-intensity stimulation, and yellow is an optogenetic inhibitory high-intensity stimulation.*

Continuing, for CMRO_2_ it is instead the pyramidal cells and the astrocytes that dominate the signal (Figure S1B). Only in experiment 4, an inhibitory low-intensity stimulation, is the relative contribution of the NO interneurons greater. At first glance, this is surprising, as the pyramidal cells and glial cells are the metabolic-draining cell types. However, when looking closer at this specific experiment one notices that this specific stimulation yields a change of CMRO_2_ in the order of magnitude of 10^-6^. In other words, this inhibitory low-intensity stimulation does not result in a relevant increase of the CMRO_2_ and therefore the contribution of the NO interneurons reached ~60% as their small metabolic contribution becomes significant compared to the small order of magnitude. As such, this experiment could be discarded as there is no relevant increase in metabolic activity.

Thirdly, we look at the relative contribution to the HbR response. This measure, like the BOLD signal, has a vascular component and a metabolic component and is a balance of the cerebral blood flow (CBF) blood highly saturated with blood and the metabolic load consuming oxygen. As we see in Figure S2, for most of the experiments the NO interneurons have the highest contribution. This is attributed to the neurovascular regulation providing a stronger vascular response relative to the metabolic need. As such, the CBF response generally brings in more oxygen than consumed and since the NO interneurons dominate vascular response (Figure S1A), they have a strong influence on the HbR response. In the experiments where NO interneurons do not have the highest relative contribution, experiments 1-3, we observe a relatively weak vascular response. This results in the cell type having the highest metabolic load, the astrocytes for these three experiments (Figure S1B), consume more oxygen than replenished by the CBF. In the case of these three experiments, we have two sensory stimulations and one optogenetic excitatory stimulation. As these stimulation types yield weak vascular responses, this behaviour is not surprising.

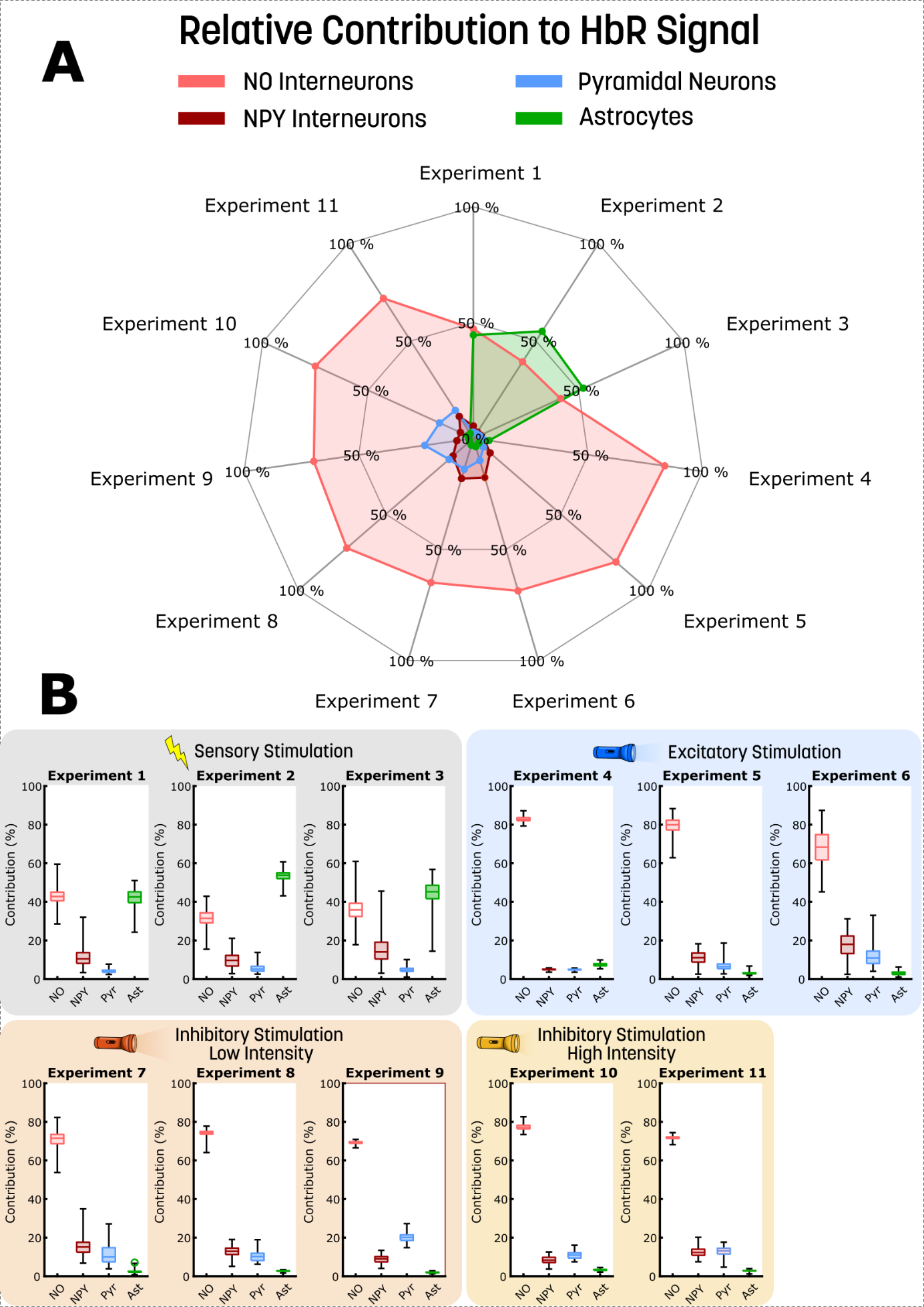

***Figure S2: The contribution of different neuronal populations to the HbR signal. A)*** *The contribution of four different neuronal populations, nitric oxide (NO) interneurons (beige), neuropeptide Y (NPY) interneurons (red), pyramidal neurons (Pyr, light blue), and glia cells (Ast, green), to the deoxygenated haemoglobin (HbR) are presented for 11 different experiments.* ***B)*** *Boxplots detailing the contribution of the four neuronal populations (NO beige, NPY red, Pyr light blue, Ast green), with uncertainty, for the 11 experiments presented in A. The background of each experiment indicates the type of stimulation that generated these behaviours: grey is a sensory stimulation, blue is an optogenetic excitatory stimulation, orange is an optogenetic inhibitory low-intensity stimulation, and yellow is an optogenetic inhibitory high-intensity stimulation.*

As the NO interneurons have the highest relative contribution to the BOLD signal, see Figure 2, it is not surprising to see that the HbR also is dominated by the NO interneurons. Generally, this analysis showcases the dominance of the vascular response in the neurovascular coupling (NVC). Furthermore, we can also observe that even the sensory and excitatory stimulation types are dependent on the inhibitory neurons in the following neurovascular regulation.

### 2 Qualitative investigations of the double-peak response attributed to SOM releasing interneurons

The double-peak dynamic, seen in Figure 7A, is generated by a slow recovery of neuronal activity following the end of stimulation. The OG stimulation, $u_{4}$, increases the neuronal activity of the somatostatin (SOM) interneurons, which in turn releases GABA. The GABA inhibits the activity of the other neuron types, pyramidal cells, NO-expressing interneurons, and NPY-expressing interneurons, for the duration of the stimulation. Figure S3A shows neuronal activity, normalized against its maximum value, meaning that 0 indicates basal activity, 1 indicates maximum activity, and -1 indicates lowest activity. After the 16 s stimulation has ceased some different behaviours are observed. The activity of SOM interneurons slowly decreases, not reaching basal levels (Figure S3A, ruby). The activity of pyramidal cells (Figure S3A, blue) and NO interneurons (Figure 7A, pink) increases above their baseline values, before returning to baseline. The NPY interneurons (Figure S3A, crimson) recovery towards baseline is slightly slower. These dynamics are propagated to the release of vasoactive substances (Figure S3B). The increased release of NO (purple) and NPY (dark red), due to the SOM interneurons, represents the biggest contribution to the vasculature. The reduced levels of PGE_2_ (Figure S3B, blue) and NPY from the NPY interneurons (Figure S3B, red) released, have a minor contribution to the vasoactive effect. Lastly, the contribution NO from the NO interneurons is neglectable (Figure S3B, pink). The slight timing difference of NO (purple) and NPY (dark red), from the SOM interneurons, produces the initial peak in the total vasoactive effect, and the difference in absolute amplitude results in the plateau before return to baseline (Figure S3C, grey line). Adding the contribution of PGE_2_ (Figure S3C, blue) gives a more pronounced initial peak due to the slower-decreasing vasodilation caused by a lack of PGE_2_. Additionally, a small secondary peak (Figure S3C, dark grey line) is observed as PGE_2_ levels recover post-stimulation. Finally, adding the NPY contribution (Figure S3C, red), the second peak is fully produced (Figure S3C, black line). Thus, the second peak behaviour is produced by the recovery of activity in the non-stimulated neurons. The reduction of NPY causes the main contribution to the second peak, as the reduction of the vasoconstricting NPY has a dilatory effect. The initial reduction of PGE_2_, followed by the small overshoot before returning to baseline, makes the second peak more pronounced.

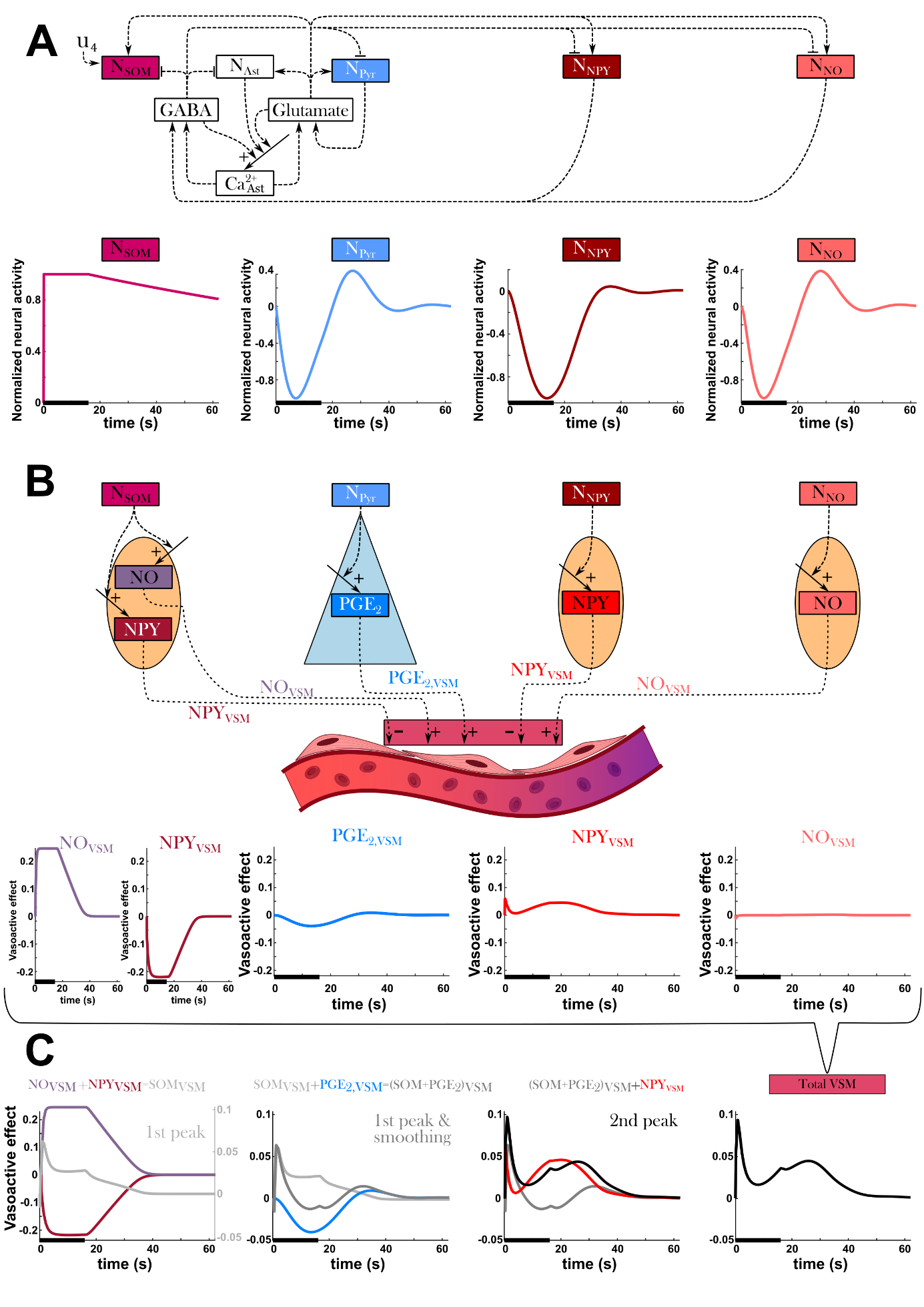
***Figure S3: The delayed dilation of somatostatin-releasing interneurons is caused by different neuronal recovery speeds post-stimulation. A)*** *A simplified overview of neural activity part of the model, including the extension of the somatostatin-releasing interneurons (SOM), for a 16 s long stimulation, u_4_, applied to the SOM neurons. As the SOM neurons (ruby) are stimulated, GABA is produced which inhibits the other neurons which are not externally stimulated. This is further highlighted by the plots of the normalized neural activity for SOM (ruby), pyramidal (blue), NPY expressing (crimson), and NO expressing (light red) neurons. For these plots, the x-axis represents time in seconds.* ***B)*** *A simplified model overview of how the neural activity propagates to the vasoactive signalling substances. NO and PGE2 dilate the vasculature, while NPY contracts. For the same 16 s long stimulation the vasoactive effect of NO (purple) and NPY (dark red), released by SOM neurons, PGE2 (blue), released by pyramidal neurons, NPY (red), released by NPY interneurons, and NO (light red), released by NO interneurons, are shown.* ***C)*** *By sequentially adding the individual vasoactive effects together, we can identify the contribution of the different substances. NO and NPY released by the SOM neurons produces the initial peak and a plateau that returns to the baseline (grey line). Adding PGE_2_ to the grey line transforms the plateau to the initial part of the second peak (dark grey line). Adding NPY released from the NPY interneurons further accentuates the initial peak and raises the amplitude of the second peak. NO released from the NO interneurons exhibit a weak impact on the total vasoactive response.*

### 3 Compilation of all experimental protocol variables

Table S1 details the experimental protocol parameters across the 17 experiments analysed in this study. The colours represent the different stimulation types: grey is sensory stimulation, light blue is optogenetic excitatory stimulation, orange is optogenetic inhibitory low-intensity stimulation, yellow is optogenetic inhibitory high-intensity stimulation, and red is optogenetic SOM stimulation. By comparing the experiments sharing the colour, and thus the stimulation type, one can observe that the protocol parameters differ greatly. For instance, the stimulation duration and frequency differ, but also factors like the anaesthesia dosing and type of agents differ across the experiments. These are attributing factors to the fact that these neurovascular systems are difficult to incorporate into a consensus view, as the effects observed are derived from different experimental protocol parameters.

In theory, one could consider the total light exposure to be the driving factor. One can calculate the total amount of light applied over the stimulation duration by multiplying the pulse frequency with the pulse width. This product gives the total light exposure for each second of the stimulation period (f*pw=amount of light/s). Notably, all optogenetic stimulation has a light exposure of 150-200 ms per second. This indicates that the configuration of delivering the light has a stronger influence compared to the total light exposure. Please see supplementary Table S1 for details.

### 4 Description of the BOLD contribution calculations

Using the formulated model, the strategy to extract the proportional contributions to the BOLD signal is described herein. Firstly, the vascular contributions are different for each neuron type. Notably, the stimulation effect on the arterioles (V1) does not scale linearly with the vascular smooth muscle (vsm) effect from the neurons. Additionally, the vsm stimulation effect can alter between having a positive and negative effect (change from basal effect). Therefore, the neuronal effect on the vasculature was estimated making use of the known volume values and fractional vsm effect, see equation S1-S4.

$$N_{frac}(t) =\frac{|vsm_{N}(t)|}{vsm_{tot}(t)}$$

$$vsm_{tot}\left( t \right)=\sum_{1}^{N} \left| vsm_{N}\left( t \right) \right|+\epsilon$$

$$N=\{NO, NPY, Pyr\}$$

(S1)

Here, the fractions at each time point (t) for NO, NPY, and Pyr are estimated from the respective absolute vsm effect over the total absolute vsm effect. $\epsilon=1e^{-10}$ avoids zero division. This gives us an absolute fraction of the effect from each neuron. However, as the effect can be negative, we cannot directly apply these fractions to the derivates. We can correct this possibility by distributing the negative effect over the positive effects, to preserve the volumes.

$$M_{N}=vsm_{N}\left( t \right)<0$$

$$N_{N}=vsm_{N}\left( t \right)\geq0$$

$$N=\{NO, NPY, Pyr\}$$

(S2)

Here, M_N_ indicates where the vsm effective is negative and N_N_ indicates where they are greater or equal than zero. With these Booleans, we can formulate the final fractions that correct for the sign and lost amount with the earlier simplification of the absolute fractions.

$$frac_{NO}\left( t \right)=\frac{vsm_{NO}(t)}{vsm_{tot}(t)}+\left( M_{NPY}*\left( \frac{NO_{frac}\left( t \right)}{NO_{frac}\left( t \right)+N_{Pyr}*Pyr_{frac}\left( t \right)}*2*NPY_{frac}\left( t \right) \right)+ M_{Pyr}*\left( \frac{NO_{frac}\left( t \right)}{NO_{frac}\left( t \right)+N_{NPY}*NPY_{frac}(t)}*2*Pyr_{frac}(t) \right) \right)*N_{NO}$$

$$frac_{NPY}\left( t \right)=\frac{vsm_{NPY}\left( t \right)}{vsm_{tot}\left( t \right)}+\left( M_{NO}*\left( \frac{N{PY}_{frac}\left( t \right)}{N{PY}_{frac}\left( t \right)+N_{Pyr}*Pyr_{frac}\left( t \right)}*2*NO_{frac}\left( t \right) \right)+ M_{Pyr}*\left( \frac{N{PY}_{frac}\left( t \right)}{N{PY}_{frac}\left( t \right)+N_{NO}*NO_{frac}\left( t \right)}*2*Pyr_{frac}\left( t \right) \right) \right)*N_{NPY}$$

$$frac_{Pyr}\left( t \right)=\frac{vsm_{Pyr}(t)}{vsm_{tot}(t)}+\left( M_{NO}*\left( \frac{{Pyr}_{frac}\left( t \right)}{{Pyr}_{frac}\left( t \right)+N_{NPY}*{NPY}_{frac}\left( t \right)}*2*{NO}_{frac}\left( t \right) \right)+ M_{NPY}*\left( \frac{{Pyr}_{frac}\left( t \right)}{{Pyr}_{frac}\left( t \right)+N_{NO}*{NO}_{frac}(t)}*2*{NPY}_{frac}(t) \right) \right)*N_{Pyr}$$

(S3)

Now the respective volumes can be calculated from the original volume values.

$$V1_{N}=V1\left( 0 \right)+frac_{N}\left( t \right)*\left( V1\left( t \right)-V1\left( 0 \right) \right)$$

$$V2_{N}=V2\left( 0 \right)+frac_{N}\left( t \right)*\left( V2\left( t \right)-V2\left( 0 \right) \right)$$

$$V3_{N}=V3\left( 0 \right)+frac_{N}\left( t \right)*\left( V3\left( t \right)-V3\left( 0 \right) \right)$$

$$N=\{NO, NPY, Pyr\}$$

(S4)

Continuing, we need to estimate the effect on deoxyhaemoglobin (HbR) from each neuron type. First, we establish the effect of the active cerebral metabolic rate of oxygen (CMRO_2_), above the resting metabolism, of the different neurons (NO, NPY, Pyr, Ast).

$$CMRO_{2,N}\left( t \right)=CMRO_{2}\left( 0 \right)*k_{CMRO_{2,N}}*\left( Ca_{N}\left( t \right)-Ca_{N}\left( 0 \right) \right)$$

$$N=\{NO, NPY, Pyr, Ast\}$$

(S5)

As the consumed amount of oxygen, jO_2_, is calculated from the metabolic activity of all neurons, we need to calculate the amount of oxygen that would not have been consumed if only one neuron type was active. We can do this via the active CMRO_2_ contributions established in equation S5. As the basal metabolism is included in jO_2_ we must account for the whole CMRO_2_ effect.

$$jO_{2,i}NO\left( t \right)=jO_{2,i}\left( t \right)*\left( \frac{CMRO_{2,NPY}(t)+CMRO_{2,Pyr}(t)+CMRO_{2,Ast}(t)}{CMRO_{2}\left( t \right)} \right)$$

$$jO_{2,i}NPY\left( t \right)=jO_{2,i}\left( t \right)*\left( \frac{CMRO_{2,NO}(t)+CMRO_{2,Pyr}(t)+CMRO_{2,Ast}\left( t \right)}{CMRO_{2}\left( t \right)} \right)$$

$$jO_{2,i}Pyr\left( t \right)=jO_{2,i}\left( t \right)*\left( \frac{CMRO_{2,NO}(t)+CMRO_{2,NPY}(t)+CMRO_{2,Ast}(t)}{CMRO_{2}\left( t \right)} \right)$$

$$jO_{2,i}Ast\left( t \right)=jO_{2,i}\left( t \right)*\left( \frac{CMRO_{2,NO}(t)+CMRO_{2,NPY}(t)+CMRO_{2,Pyr}\left( t \right)}{CMRO_{2}\left( t \right)} \right)$$

$$i=\{1,2,3\}$$

(S6)

Now we can estimate the oxygen amount for each neuron pathway with corrected metabolic effect.

$$nO_{2,i}N_{met}\left( t \right)=nO_{2,i}\left( t \right)+jO_{2,i}N\left( t \right)$$

$$N=\{NO, NPY, Pyr, Ast\}$$

$$i=\{1,2,3\}$$

(S7)

Further, we now also estimate the full oxygen amount accounting for the vascular changes from the vsm.

$$nO_{2,i}N\left( t \right)=nO_{2,i}N_{met}\left( 0 \right)+frac_{N}\left( t \right)*(nO_{2,i}N_{met}\left( t \right)-nO_{2,i}N_{met}(0)$$

$$N=\{NO, NPY, Pyr, Ast\}$$

$$i=\left\{ 1,2,3 \right\}$$

$$frac_{Ast}=0$$

(S8)

We can now calculate the oxygen concentrations.

$$cO_{2,i,i+1}N\left( t \right)=\frac{nO_{2,i}N\left( t \right)}{V_{i}N\left( t \right)}$$

$$N=\{NO, NPY, Pyr, Ast\}$$

$$i=\left\{ 1,2,3 \right\}$$

$$V_{i}Ast\left( t \right)=V_{i}\left( t \right)$$

(S9)

Thereafter the oxygen saturations.

$$S_{i}O_{2}N\left( t \right)=min(0.95,\frac{cO_{2,i,i+1}\left( t \right)+cO_{2,i+1,i+2}\left( t \right)}{2*cO_{2,max}}$$

$$cO_{2,max}=9.26$$

$$N=\left\{ NO, NPY, Pyr, Ast \right\}$$

$$i=\left\{ 1,2,3 \right\}$$

(S10)

And then the HbR contents.

$$HbR_{i,N}\left( t \right)=Vi_{N}\left( t \right)*\left( 1-S_{i}O_{2}N\left( t \right) \right)$$

$$HbR_{N}\left( t \right)=\sum_{i=1}^{i} HbR_{i,N}\left( t \right)$$

$$N=\{NO, NPY, Pyr, Ast\}$$

$$i=\left\{ 1,2,3 \right\}$$

(S11)

Since there is a physiological cap of oxygen saturation (95%), we must correct for the lost HbR due to the capped saturation. We distribute this HbR based on the metabolic activity of the neurons.

$$HbR_{missing}\left( t \right)= \sum_{i}^{N} HbR_{N}\left( t \right)-HbR_{N}\left( 0 \right)$$

$$CMRO_{2,tot}(t)=\sum_{1}^{N} \left| CMRO_{2,N}\left( t \right) \right|+\epsilon$$

$$HbR_{N}\left( t \right)=HbR_{N}\left( t \right)+\frac{\left| CMRO_{2,N}\left( t \right) \right|}{CMRO_{2,tot}\left( t \right)}*HbR_{missing}\left( t \right)$$

$$N=\{NO, NPY, Pyr, Ast\}$$

(S12)

Now we can finally calculate the OIS-BOLD signal from the different neurons.

$$CBV_{N}\left( t \right)=\sum_{i=1}^{3} Vi_{N}\left( t \right)$$

$$BOLD_{N}(t)=\exp\left( -k_{y,BOLD}*(HbR_{N}\left( t \right)-HbR_{N}\left( 0 \right)-k_{y,BOLD2}*(CBV_{N}\left( t \right)-CBV_{N}(0) \right)$$

$$N=\{NO, NPY, Pyr, Ast\}$$

(S13)

Here k_y, BOLD_ and k_y, BOLD2_ are parameters. We then use the area of the signal to translate from the time series to the percentual contributions.

$$BOLD_{N,area}=trapz\left( \left| BOLD_{N}\left( t \right) \right|-1 \right)$$

$$BOLD_{total}=\sum_{1}^{N} BOLD_{N,area}$$

$$BOLD_{N,contribution}=100*\frac{BOLD_{N,area}}{BOLD_{total}}$$

$$N=\{NO, NPY, Pyr, Ast\}$$

(S14)

### 5 Full model description

The model interaction graph is depicted in Figure 8 and describes all reactions and interactions of the model. In practice, the presented model builds on the work of Sten *et al.* (1), which included submodules from various previous studies (2–6). Note that the full model description is given, including the somatostatin (SOM) module. To get the more condensed model version that excludes the SOM module, simply remove these equations and effects. Following, all model equations will be presented. The stimulus $u$ is given by the following equation:

$u_{i}=\left\{ \begin{aligned} 1 t_{on}\leq t\leq t_{off} \\ 0 otherwise \end{aligned} \right.$	(S15)

where $t_{\mathrm{on}}$, $t_{\mathrm{off}}$ are the times when the signal goes on and off, respectively.

#### 5.1 Presynaptic activity and calcium influx

The neuronal activity $N_{i}$ states are represented such as:

$\frac{d}{dt}{[N}_{NO}]=k_{u1}*u_{1}*\max\left( N_{max,NO}-\left[ N_{NO} \right],0 \right)+k_{PF,1}*\left[ Glut \right]+k_{IN,1}*\left[ GABA \right]-sinkN_{NO}*[N_{NO}]$	(S16a)

$\frac{d}{dt}{[N}_{NPY}]=k_{u2}*u_{2}*\max\left( N_{max,NPY}-{[N}_{NPY}],0 \right)+k_{PF,2}*[Glut]+k_{IN,2}*[GABA]-sinkN_{NPY}*{[N}_{NPY}]$	(S16b)

$\frac{d}{dt}{[N}_{Pyr}]=k_{u3}*u_{3}*\max\left( N_{max,Pyr}-\left[ N_{Pyr} \right],0 \right)+k_{PF,3}*\left[ Glut \right]+k_{IN,3}*\left[ GABA \right]-sinkN_{Pyr}*[N_{Pyr}]$	(S16c)

$\frac{d}{dt}{[N}_{Ast}]=k_{PF,4}*[Glut]+k_{IN,4}*[GABA]-sinkN_{Ast}*{[N}_{Ast}]$	(S16d)

$\frac{d}{dt}{[N}_{SOM}]=k_{u4}*u_{4}*\max\left( N_{max,SOM}-\left[ N_{SOM} \right],0 \right)+k_{PF,5}*\left[ Glut \right]+k_{IN,5}*\left[ GABA \right]-sinkN_{SOM}*[N_{SOM}]$	(S16e)

Where $u_{i}$, and $i=\{1, 2, 3, 4\}$, is the stimulation function; $N_{max,j}$, and $j=\{NO, NPY, Pyr, Ast, SOM\}$, is the parameter determining the saturation in the stimulating effect i.e., the stimulation term becomes 0 when $N_{j}>N_{max,j}$; glutamate and GABA are the changes in glutamate and GABA levels respectively; $k_{u,i}$,$k_{PF,i}$, $k_{IN,i}$, and $sinkN_{j}$ are scaling parameters scaling the effects of the stimulation, the neuronal activation caused by glutamate, the neuronal inhibition caused by GABA, and the general return to baseline neuronal activity, respectively. The levels of glutamate and GABA are implemented as

$$\frac{d}{dt}\left[ GABA \right]=k_{GABA,NO}*\left[ Ca_{NO}^{2+} \right]+k_{GABA,NPY}*\left[ Ca_{NPY}^{2+} \right]+k_{GABA,Ast}*\left[ Ca_{Ast}^{2+} \right]+k_{GABA,SOM}*\left[ Ca_{SOM}^{2+} \right]-sink_{GABA}*[GABA]$$

(S17)

$$\frac{d}{dt}\left[ Glut \right]=k_{Glut,Pyr}*\left[ Ca_{Pyr}^{2+} \right]+k_{Glut,Ast}*\left[ Ca_{Ast}^{2+} \right]- sink_{Glut}*[Glut]$$

(S18)

Where $k_{Glut,j}$and $k_{GABA,j}$ are kinetic rate parameters determining the release of glutamate and GABA from the respective neuron populations; $Ca_{j}^{2+}$ are the $Ca^{2+}$ levels for the respective neuron populations; $sink_{j}$ is a parameter determining the rate at which the glutamate and GABA levels return to baseline.

The differential equations for the $Ca_{j}^{2+}$ states are formulated as:

$$\frac{d}{dt}\left[ Ca_{NO}^{2+} \right]={ln(1+exp[k}_{Ca}+k_{Ca,NO}*{[N}_{NO}]])-{sinkCa}_{NO}*{[Ca}_{NO}^{2+}]$$

(S19a)

$$\frac{d}{dt}\left[ Ca_{NPY}^{2+} \right]={ln(1+exp[k}_{Ca}+k_{Ca,NPU}*[N_{NPY}]])-{sinkCa}_{NPY}*{[Ca}_{NPY}^{2+}]$$

(S19b)

$$\frac{d}{dt}\left[ Ca_{Pyr}^{2+} \right]={ln(1+exp[k}_{Ca}+k_{Ca,Pyr}*{[N}_{Pyr}]])-{sinkCa}_{Pyr}*{[Ca}_{Pyr}^{2+}]$$

(S19c)

$$\frac{d}{dt}[Ca_{Ast}^{2+}]={ln(1+exp[k}_{Ca}+k_{Ca,Ast}*{[N}_{ASt}]])+kPFCa*[Glut]+kPF2Ca*[GABA]-{sinkCa}_{Ast}*[{Ca}_{Ast}^{2+}]$$

(S19d)

$$\frac{d}{dt}[Ca_{SOm}^{2+}]={ln(1+exp[k}_{Ca}+k_{Ca,SOM}*{[N}_{SOM}]])-{sinkCa}_{SOM}*[{Ca}_{SOM}^{2+}]$$

(S19e)

where $Ca_{j}^{2+}$is the collective level of $Ca^{2+}$for neuron type $j$; $k_{Ca}$ and $k_{Ca,j}$ are general and neuron-type-specific scaling parameter respectively; $N_{j}$ is the neuronal activity of neuron type $j$; and $sinkCa_{j}^{2+}$ is a collective term for the reduction in the $Ca^{2+}$levels.

#### 5.2 Pyramidal neuron signalling

The rise in intracellular Ca^2+^ levels in pyramidal neurons activates phospholipases which metabolize, through intermediary enzymatic steps, membrane phospholipids into intracellular arachidonic acid (AA) (7–9). This is described by:

$\frac{d\left[ AA \right]}{dt}=k_{PL}\left[ Ca_{Pyr}^{2+} \right]-\frac{k_{COX}\left[ AA \right]}{K_{M,COX}+[AA]}$	(S20)

where $k_{\mathrm{PL}}$ and $k_{\mathrm{COX}}$ are kinetic rate parameters. In pyramidal neurons, AA is metabolized into prostaglandin E_2_ (PGE_2_) through a cyclooxygenase-2 (COX-2) and PGE synthase rate-limiting reaction (9,10). PGE_2_ evokes vasodilation through the activation of EP2 and EP4 receptors expressed on the surface of vascular smooth muscle cells (VSM) cells (9,11). In the model, this mechanism is described as

$\frac{d\left[ PGE_{2} \right]}{dt}=\frac{k_{COX}\left[ AA \right]}{K_{m,COX+[AA]}}-k_{PGE_{2}}[PGE_{2}]$	(S21a)

$\frac{d\left[ PGE_{2,vsm} \right]}{dt}=k_{PGE_{2}}\left[ PGE_{2} \right]-sink_{PGE_{2}}[PGE_{2,vsm}]$	(S21b)

Where $\mathrm{PGE}_{2, vsm}$ represents PGE_2_ acting on the VSM cells, and $k_{PGE2}$ and ${sink}_{PGE2}$ are kinetic rate parameters.

#### 5.3 GABAergic interneuron signalling

The rise in intracellular Ca^2+^ levels in GABAergic interneurons evokes the release of different vasoactive messengers and substances. For instance, NO has previously been shown to be a potent vasodilator at the level of arteries and arterioles in both *in vitro* and *in vivo* studies (11–14). NO is released by specific NO-interneurons through a Ca^2+^-dependent nitric oxide synthase (NOS) rate-limiting reaction. Here, we represent this process as

$\frac{d\left[ NO \right]}{dt}=k_{NOS}\left[ Ca_{NO}^{2+} \right]-k_{NO}*\frac{\left[ NO \right]}{kM_{NO}+[NO]}$	(S22a)

$\frac{d\left[ SOM_{NO} \right]}{dt}=k_{NOS2}\left[ Ca_{SOM}^{2+} \right]-k_{NO2}*\frac{\left[ SOM_{NO} \right]}{kM_{NO2}+[SOM_{NO}]}$	(S22b)

$\frac{d\left[ NO_{vsm} \right]}{dt}=k_{NO}*\frac{\left[ NO \right]}{kM_{NO}+\left[ NO \right]}+k_{NO2}*\frac{\left[ SOM_{NO} \right]}{kM_{NO2}+[SOM_{NO}]}-sink_{NO}[NO_{vsm}]$	(S22c)

where $\mathrm{NO}_{\mathrm{vsm}}$ represents NO acting on the VSM cells, and $k_{\mathrm{NOS}}$, $k_{NOS2}$, and ${sink}_{\mathrm{NO}}$ are kinetic rate parameters. The conversion between intracellular NO and NO in the VSM, $NO_{vsm}$, is governed by Michaelis-Menten kinetics, where ${kM}_{\mathrm{NO}}$ and ${kM}_{NO2}$ are the Michaelis constant the concentration of the substrate at which half max reaction rate is achieved), $k_{\mathrm{NO}}$ and $k_{NO2}$ is the maximum reaction rate. Here, ${sink}_{\mathrm{NO}}>1 s^{-1}$ following experimental work by (15).

Another potent vasoactive messenger is NPY. NPY is released by specific NPY-producing subtypes of GABAergic interneurons and has previously been shown to induce vasoconstriction of vessels *in vitro* (11,16) and *in vivo* (17). NPY binds to the NPY receptor Y1, a G_αi_-protein coupled receptor expressed on the surface of VSM cells enwrapping arteries and arterioles (18–20). Activation of this receptor inhibits the synthesis of adenosine monophosphate (cAMP) and increases the intracellular Ca^2+^ (21), leading to VSM contraction and vasoconstriction. In the model, these mechanisms are represented as:

$\frac{d\left[ NPY \right]}{dt}=k_{NPY}\left[ Ca_{NPY}^{2+} \right]-\frac{V_{max}\left[ NPY \right]}{{kM}_{NPY}+\left[ NpY \right]}$	(S23a)

$\frac{d\left[ SOM_{NPY} \right]}{dt}=k_{NPY2}\left[ Ca_{SOM}^{2+} \right]-\frac{V_{max2}\left[ SOM_{NPY} \right]}{{kM}_{NPY2}+\left[ SOM_{NPY} \right]}$	(S23b)

$\frac{d\left[ NPY_{vsm} \right]}{dt}=\frac{V_{max}\left[ NPY \right]}{{kM}_{NPY}+\left[ NpY \right]}+\frac{V_{max2}\left[ SOM_{NPY} \right]}{{kM}_{NPY}+\left[ SOM_{NPY} \right]}-sink_{NPY}[NPY_{vsm}]$	(S23c)

where $\mathrm{NPY}_{\mathrm{vsm}}$ represents NPY acting on the VSM cells, and $k_{\mathrm{NPY}}$, $k_{NPY2}$, and ${sink}_{\mathrm{NPY}}$ are kinetic rate parameters. The conversion between intracellular NPY and NPY in the VSM, $\mathrm{NPY}_{\mathrm{vsm}}$, is governed by Michaelis-Menten kinetics (22,23), where ${kM}_{\mathrm{NPY}}$ and ${kM}_{NPY2}$ is the Michaelis constant (and $V_{\max}$and $V_{max2}$is the maximal reaction rate.

The expression of the different vasoactive substances is scaled with a parameter, $ky_{h}, h=\{NO, PGE2, NPY\}$, and summarized to a total vascular influence $G$:

$G=k_{y,NO}\left( \left[ NO_{vsm} \right]-\left[ NO_{vsm,0} \right] \right)+k_{y,PGE_{2}}\left( \left[ PGE_{2,vsm} \right]-\left[ PGE_{2 vsm,0} \right] \right)- k_{y,NPY}(\left[ N{PY}_{vsm} \right]-[N{PY}_{vsm,0}]$) (S24)

#### 5.4 Cerebrovascular dynamics

We describe the dynamics of three cerebrovascular compartments, corresponding to arteries/arterioles ($a$), capillaries ($c$), and veins/venules ($v$), respectively. The compartments are represented by an electrical circuit analogy, as originally presented in the work by Barrett *et al.* (3–5)*.* We use the derivative work of (24) and only present the fully derived equations to improve the clarity for the reader. We refer the reader to the original articles for an in-depth description of the equations (3–5,24). The following equations describe the volume change for each respective cerebrovascular compartment, $V_{i}, with i=\{a, c, v\}$.

$\frac{d[V_{a}]}{dt}=\frac{1}{k_{vis,a}}\left( \frac{K_{a}-\frac{[V_{a}]}{V_{a,0}}}{K_{a}-1}+G-\frac{2}{C_{a,0}}\frac{{[V}_{a}]}{\left( r_{1}+r_{2} \right)*f_{1}+\left( r_{2}+r_{3} \right)*f_{2+r_{3}f_{3}}} \right)$	(S25a)

$\frac{d{[V}_{c]}}{dt}=\frac{1}{k_{vis,c}}\left( \frac{K_{c}-\frac{[V_{c}]}{V_{c,0}}}{K_{c}-1}-\frac{2}{C_{c,0}}\frac{{[V}_{c]}}{\left( r_{2}+r_{3} \right)*f_{2+r_{3}f_{3}}} \right)$	(S25b)

$\frac{d{[V}_{v}]}{dt}=\frac{1}{k_{vis,v}}\left( \frac{K_{v}-\frac{[V_{v}]}{V_{v,0}}}{K_{v}-1}-\frac{2}{C_{v,0}}\frac{{[V}_{v}]}{r_{3}f_{3}} \right)$	(S25c)

Here, $K$ are stiffness coefficients; $k_{vis,i}$ are viscoelastic parameters; $C_{i}$ represents the compliance of the vessel; $R$ represents the vessel resistance; $f$ represents the flow of blood between the cerebrovascular compartments; the baseline value is indicated by the subscript $0$, and finally, $G$is the vasoactive function translating the actions of the vasoactive arms (Eq. S24) into hemodynamic changes.

In short, the arterial compartment is actively regulated by the vasoactive arms (Eq. S21b, S22c, and S23c), and the evoked hemodynamic changes in the arterial vessels are propagated through the capillary and venous compartments.

Using conservation of mass, the rate at which the blood volume changes in a compartment is given by the difference between the in- and outflow of blood:

$\frac{d{[V}_{i}]}{dt}=f_{i-1}-f_{i}$	(S26)

Restructuring the equation gives the following relationship:

$f_{i}=f_{i-1}-\frac{d[V_{i}]}{dt}$	(S27)

Next, the pressure drop over a compartment is given by:

$\Delta p_{i}\left( t \right)=\frac{1}{2}\sum_{i=1} r_{i}(f_{i-1}+f_{i})$	(S28)

Which is subject to the pressure boundary conditions:

$\sum_{i=1}^{3} \Delta p_{i}=\Delta p_{r}=1$	(S29)

Using these conditions, we can solve for the inflow of blood into the first compartment $f_{0}$:

$f_{0}=\frac{2-(\left( r_{1}+r_{2} \right)f_{1}+\left( r_{2}+r_{3} \right)f_{2}+r_{3}f_{3})}{r_{1}}$	(S30)

The equations above form a DAE system, comprised of three volumes and four flows. To further simplify, the scales of the variables are normalized to:

$\left[ \begin{aligned} \sum_{i=a}^{v} V_{i,0}=1 \\ f_{i,0}=1 \\ \sum_{i=a}^{v} r_{i,0}=1 \end{aligned} \right]$	(S31)

Using this, the rest of the physiological terms can be expressed as:

$p_{i,0}=\left\{ \begin{aligned} \Delta p_{r} if i=a \\ \Delta p_{r}-f_{i,0}\sum_{j=1}^{i-1} r_{j,0} if i=\{c,v\} \end{aligned} \right.$	(S32a)

$c_{i,0}=\frac{V_{i,0}}{p_{i,0}-\frac{1}{2}r_{i,0}f_{i,0}}$	(S32b)

$l_{i}=\left( r_{i,0}v_{i,0}^{2} \right)^{\frac{1}{3}}$	(S32c)

$r_{i}=\frac{l_{i}^{3}}{v_{i}^{2}}$	(S32d)

where $P$ is the pressure at the entry point to each compartment, $C$ is the compliance of the vascular vessel, $L$ is the length of the vascular segment, and $R$ is the vessel resistance to flow.

Given Equations S25–S32, the initial conditions for the cerebrovascular volumes $V_{i}$ and the vascular resistance $R_{i}$ must be specified. We set these initial conditions in accordance with the work of Barrett *et al*. (3), which based these values on the available literature, resulting in $V_{i,0}=[0.29, 0.44, 0.27]$ and $R_{i,0}=\left[ 0.74, 0.08, 0.18 \right]$.

#### 5.5 Oxygen transportation

To describe the oxygen transportation through the cerebrovascular system, we describe three vascular compartments and a cerebral tissue compartment. The amount of oxygen in these compartments is given by:

$\frac{d[n_{O_{2},a}]}{dt}=f_{0}C_{O_{2},in}-f_{1}C_{O_{2},a,c}-j_{O_{2},a}-j_{O_{2},s}$	(S33a)

$\frac{d{[n}_{O_{2},c}]}{dt}=f_{1}C_{O_{2},a,c}-f_{2}C_{O_{2},c,v}-j_{O_{2},c}$	(S33b)

$\frac{d{[n}_{O_{2},v]}}{dt}=f_{2}C_{O_{2},c,v}-f_{3}C_{O_{2},out}-j_{O_{2},v}+j_{O_{2},s}$	(S33c)

$\frac{d{[n}_{O_{2},t]}}{dt}=\sum_{i} j_{O_{2},i}-CMRO_{2}$	(S33d)

where the flow of blood $f$ transports oxygen in and out from each compartment. The permeability in the vessel wall allows for oxygen to diffuse to the tissue compartment, $jO2_{i}$, where $i=\{a,c,v\}$. An arterio-venous diffusion shunt is described by $jO2_{s}$, where oxygen moves from the arteriole to the venous compartment. Oxygen is metabolized in the tissue compartment, ${CMRO}_{2}$. The metabolism of oxygen is given by:

$CMRO_{2}=CMRO_{2,ss}*(1+\sum_{\forall j} k_{CMRO_{2},j}*Ca_{j}^{2+})$	(S34)

where ${CMRO}_{2,0}$ is the basal metabolism of oxygen, which is increased by the activity level of each neuron (see Eq. S16) scaled by the parameter $k_{CMRO_{2},j}$, where $j=\{NO, NPY, Pyr, Ast, SOM\}$.

The concentration of oxygen, $C_{O_{2},i}$ is given by the amount of oxygen, $n_{O_{2},i}$, divided by the volume of each compartment, $V_{i}$, except for the oxygen concentration that enters the arteries, $c_{O_{2},0}$, which is assumed to be constant, minus a loss of oxygen to the surrounding tissue $c_{O_{2},leak}$.

$C_{O_{2},i,i+1}=\left\{ \begin{aligned} \frac{n_{O_{2},i}}{V_{i}} if i=\{a,c,v,t\} \\ C_{O_{2},0}-C_{O_{2},leak} if i=in \end{aligned} \right.$	(S35)

Where the tissue volume fraction $V_{t}=34.8$ (4) and $c_{O_{2},leak}=0.116 mM$ (calculated using data from (4,25)).

As in the original work, by ignoring the minor fraction of oxygen directly dissolved in the blood plasma, blood oxygen concentration can be related to oxygen partial pressures through the oxygen-haemoglobin saturation curve:

$p_{O_{2},i,i+1}=p_{50}\left( \frac{C_{O_{2},max}}{C_{O_{2},i,i+1}}-1 \right)^{-\frac{1}{h}}$	(S36)

Where $p_{50}=36 mmHg$ is the oxygen partial pressure at the halfway saturation point, $cc_{O_{2},max}=9.26 mM$ is the maximum oxygen concentration in blood, and $h=2. 6$ is the Hill exponent. These values are taken from (26).

The oxygen pressure in the tissue, $p_{O_{2},t},$ is calculated using Henry’s law:

$p_{0_{2},t}=\frac{C_{O_{2},t}}{\sigma_{oxygen}}$	(S37)

Where $\sigma_{Oxygen}=1.46 \mu M/mmHg$ is the coefficient for solubility of oxygen in tissue (27).

The average pressure, $\bar{p}_{O_{2},i}$, in a compartment is obtained by averaging over the input and output pressures:

$\tilde{p}_{O_{2},i}=\frac{p_{O_{2},i-1}+p_{O_{2},i+1}}{2}$	(S38)

The diffusion of oxygen between the vessels and the tissue, $j_{O_{2},i}$, as well as between the arteries and the veins,$j_{O_{2},s}$, is driven by the difference in partial oxygen pressure:

$j_{O_{2},i}=g_{i}\left( \tilde{p}_{O_{2},i}-p_{O_{2},t} \right), i=\{a,c,v\}$	(S39a)

$j_{O_{2},s}=g_{s}(\tilde{p}_{O_{2},a}-\tilde{p}_{O_{2},v})$	(S39b)

where $g_{i} , i=\{a,v,s\}$ are rate constants. $g_{2}$ is instead given by

$g_{2}=g_{2,ss}*(1-k_{capPerm}*[f_{0}-f_{0,ss}])$	(S40)

where $g_{2,ss}$ is the permeability at baseline; $k_{capPerm}$ is a scaling parameter; $f_{0}$ is the blood flow entering the local region; and $f_{0,ss}$ is the blood flow at baseline.

We employed the identical strategy of Barrett and Suresh (4) (see supplementary material in (4)), and used the experimental $pO_{2}$ data from Vovenko (25) to calculate $g_{i}$ and ${CMRO}_{2,0}$ and estimate $C_{o_{2},leak}$ as:

$C_{o_{2},leak}=max(C_{o_{2},leak,ss}*[1-k_{leak}*(f_{0}-f_{0,ss})])$	(S41)

Where $C_{o_{2},leak}$ is the loss of O_2_ concentration due to “leakage”; $C_{o_{2},leak,ss}$ is the leakage at baseline; $k_{leak}$ is a scaling parameter; and $f_{0}$ and $f_{0,ss}$ is the blood flow and basal blood flow as described above. The $max(x,0)$function is to prevent a negative O_2_ leakage i.e., the blood entering the local region of the brain cannot have a higher O2 concentration than the systemic arteries. This leaves the parameters $k_{CMRO_{2},j}$ and $k_{leak}$ to be estimated where applicable.

The oxygen saturation of blood in the different vascular compartments, $S_{i}O_{2}$, is approximated by dividing the average oxygen concentration of each compartment with$c_{O_{2},max}$:

$S_{i}O_{2}=\frac{C_{O_{2},i-1,i}-C_{O_{2},i,i+1}}{C_{O_{2},max}}, i=\{a,c,v\}$	(S42)

Lastly, using the blood oxygen saturation $S_{i}O_{2}$ and vascular volumes $(V_{i})$, corresponding changes in oxygenated haemoglobin (HbO), deoxyhemoglobin (HbR), and total haemoglobin (HbT) are given by:

$n_{Hbt,i}={[V}_{i}]$	(S43a)

$n_{HbO,i}={[V}_{i}]*S_{i}O_{2}$	(S43b)

$n_{HbR,i}={[V}_{i}]*(1-S_{i} O_{2})$	(S43c)

#### 5.6 BOLD signal derivation

Finally, the oxygen saturation of blood in the different vascular compartments, $S_{i}O_{2}$ (Eq. S42) and the vascular volumes, $V_{i}$ (Eq. S25) are used to calculate the BOLD signal. The following derivation of the BOLD signal equations and parameter values are taken from the work of (2), and we refer the reader to the original article for a detailed view of the equations. The BOLD signal is expressed as a summation of the contributions from each vascular compartment $\{a,c,v\}$ and an extracellular compartment $\{e\}$ symbolizing the tissue:

$BOLD=H\sum_{j} S_{j}, j=\{a,c,v,e\}$	(S44a)

$H=\frac{1}{V_{e}+\sum_{k} \varepsilon_{k}\hat{V}_{k}}, k=\{a,c,v\}$	(S44b)

$\varepsilon_{(a,c,v)}=\frac{S_{\left( a,c,v \right),0}}{S_{e,0}}=\lambda\frac{e^{-TER_{2(a,c,v),0}^{*}}}{e^{-TER_{2e,0}^{*}}}$	(S44c)

Where $S_{j}$ is the intrinsic signal from each compartment; $V_{e}=1-\hat{V}_{k}$ is the extravascular volume; $\hat{V}_{k}=V_{iv,0}\sum V_{k}$ is the intravascular volume scaled by the baseline intravascular volume fraction $(V_{iv,0}=0.05$, (28)); $\varepsilon_{i}$ is the intrinsic signal ratio of blood and $\lambda$ is the intravascular to extravascular spin density ratio (we assume $\lambda=1.15$); $R_{2}^{*}$ is the transverse signal relaxation rate for the four compartments, and $TE$ is the echo time used during image acquisition in each study.

The transverse signal relaxation rate for the extravascular compartment ($R_{2e,0}^{*})$ was assumed to be $25.1s^{-1}$ (29), and was calculated for the intravascular compartments by utilizing the following quadratic expression taken from (30):

$R_{2i,0}^{*}=A^{*}+C^{*}\left( 1-S_{i}O_{2} \right)^{2}$	(S45a)

Where:

${A^{*} = 14.87Hct + 14.686 \atop C^{*} = 302.06Hct + 41.83}$	(S45b)

These equations are governed by $S_{i}O2$ (Eq. S42) and the resting haematocrit in the blood ($Hct; {Hct}_{a,v}=0.44, {Hct}_{c}=0.33$ (31,32)).

The respective signal contribution from each compartment is given by:

$S_{i}=\left\{ \begin{aligned} \varepsilon_{i}\hat{V_{i}}e^{-TE\Delta R_{2i}^{*}} if i=\{a,c,v\} \\ V_{e}e^{-TE\Delta R_{2e}^{*}} if i=e \end{aligned} \right.$	(S46)

Where $\Delta R_{2}^{*}$ (*i.e.,* the change in MR signal relaxation rate) is given by the following expressions:

$\Delta R_{2i}^{*}=C^{*}\left( \left( 1-S_{i}O_{2} \right)^{2}-\left( 1-S_{i}O_{2,0} \right)^{2} \right)$	(S47a)

$\Delta R_{2e(a,c,v)}^{*}=\left\{ \begin{aligned} \frac{4\pi}{3}\Delta_{x}Hct\gamma B_{0}\sum_{i=a,v} \left[ \hat{V_{i}}\left( \left| S_{off}O_{2}-S_{i}O_{2} \right| \right)-\hat{V_{i,0}}\left( \left| S_{off}O_{2}-S_{i}O_{2,0} \right| \right) \right] \\ 0.04\left( \Delta_{x}Hct\gamma B_{0} \right)^{2}\left[ \hat{V_{c}}\left( \left| S_{off}O_{2}-S_{c}O_{2} \right| \right)^{2}-\hat{V_{c,0}}\left( \left| S_{off}O_{2}-S_{c}O_{2,0} \right| \right) \right] \end{aligned} \right.$	(S47b)

Here, $\Delta\chi=2.64\times{10}^{-7}$ is the susceptibility of fully deoxygenated blood (33); $\gamma=2.68\times{10}^{8}$ is the gyromagnetic ratio of protons; $B_{0}$ is the magnetic field strength used during image acquisition, and finally, $S_{off}O_{2}=0.95$ is the blood saturation which gives no magnetic susceptibility difference between blood and tissue (33).

The OIS-BOLD signal was implemented in the model as an exponential expression of the relative change in HbR and CBV levels. The HbR component represents the approximation of the OIS-BOLD signal as an inversed measurement of the change in HbR. The volume-based offset was introduced to account for the light scattering effect caused by an increased blood volume i.e., an increased blood volume leads to increased light scattering which means a decrease in optical signal. This relationship is explained by the following expression:

$BOLD_{OIS}=\exp\left( -k_{BOLD}*\left( HbR-HbR_{ss} \right)-k_{BOLD2}*\left( CBV-CBV_{ss} \right) \right)$	(S48)

where $k_{BOLD}$ and $k_{BOLD2}$ are scaling parameters which are estimated from data; $HbR$ is the calculated levels of deoxygenated haemoglobin; $CBV$ is the calculated cerebral blood volume; and $HbR_{ss}$ and $CBV_{ss}$ are the baseline values of HbR and CBV, respectively.

### Supplementary Table Legends

***Table S1: Overview of the experimental protocol parameters.*** The experimental stimulation protocol parameters are detailed for the 17 experiments analysed. The table includes the study of origin, stimulation type, stimulation duration, stimulation frequency, stimulation pulse duration in milliseconds and seconds, light exposure, stimulation strength or effect, the type of mouse and the utilized mouse strain, and the dosing of the anaesthesia agents. The colours indicate the stimulation type, where: grey is sensory stimulation, light blue is optogenetic excitatory stimulation, orange is optogenetic inhibitory low-intensity stimulation, yellow is optogenetic inhibitory high-intensity stimulation, and red is optogenetic somasomatic stimulation.

***Table S2: Model parameter values:*** The model parameters are detailed in this table in log10 scale. The columns detail: the parameter name, the best estimated value found during parameter estimation, the lower bound used during parameter estimation, the upper bound used during parameter estimation, the lowest value found for the confidence interval, and the highest value found for the confidence interval. Repeated parameters are named with an abbreviation to indicate which study and experiment they belong to.

***Table S3: SOM model parameter values:*** The model parameters are detailed in this table in log10 scale. The columns detail: the parameter name, the best estimated value found during parameter estimation, the lower bound used during parameter estimation, the upper bound used during parameter estimation, the lowest value found for the confidence interval, and the highest value found for the confidence interval. Repeated parameters are named with an abbreviation to indicate which experiment they belong to.
